# Thermal and non-thermal stress conditions activate the *Plasmodium falciparum* AP2-HS-dependent heat-shock response

**DOI:** 10.1101/2025.11.24.690237

**Authors:** Neus Ràfols, Marc Chillarón-Adán, Elisabet Tintó-Font, Alfred Cortés

## Abstract

To preserve proteome homeostasis and survive at higher-than-optimal temperatures, organisms have evolved the conserved heat shock (HS) response (HSR), characterised by increased expression of specific chaperone-encoding genes. In the human blood, malaria parasites are frequently exposed to elevated temperatures associated with host fever episodes. The protective HSR of *Plasmodium falciparum*, the parasite that produces the vast majority of malaria clinical cases and deaths, is regulated by the transcription factor AP2-HS. Here, we systematically investigated the conditions that trigger the AP2-HS-dependent HSR and found that even mild HS conditions that do not compromise parasite viability can activate this response. Similar to other organisms, activation of the HSR in *P. falciparum* is rapid, as it was observed after an only 10 min HS. Artemisinin (ART), a drug that produces proteome damage, also triggered the HSR, indicating that activation of the malarial HSR is not restricted to thermal stress. The AP2-HS-dependent HSR can be activated in all asexual blood stages, with the exception of very young rings, but not in intermediate or mature gametocyte stages. Accordingly, these gametocyte stages are highly sensitive to HS. Since mature gametocytes are the only stage that can mediate human-to-mosquito transmission, these results suggest that malaria patients with high fever may become transiently uninfective.

## INTRODUCTION

Malaria continues to be one of the most important human infectious diseases, with ∼260 million cases and ∼600,000 deaths reported every year. *Plasmodium falciparum* causes the most severe form of human malaria and accounts for the vast majority of clinical cases and deaths [1]. Humans are infected through the bite of an infected *Anopheles* female mosquito. After exponential multiplication in the liver, parasites at the merozoite stage are released and start the intraerythrocytic developmental cycle (IDC). Merozoites invade red blood cells (RBCs) and progress through the ring, trophozoite and schizont stages, the latter characterised by nuclear division that leads to the formation of up to 32 new merozoites. Approximately 48 h after RBC invasion, schizonts burst, releasing merozoites that invade new RBCs and start the cycle again. At each replicative cycle, a small percentage of the parasites convert into sexual forms called gametocytes, which sequester in the bone marrow for ∼10 days until they re-enter the peripheral circulation [2]. These mature male or female gametocytes are the only stage that can infect a mosquito during a bloodmeal, therefore playing an essential role for transmission.

Cyclical fever is the hallmark of clinical malaria. Malaria fever peaks typically last a few hours, with high temperature spikes of up to 41 °C that usually last ∼1 h. In *P. falciparum* infections, fever peaks typically show a periodicity of ∼48 h (tertian fever) and coincide with schizont rupture at the end of each round of the IDC [3–5]. The release of malaria ‘toxins’ during schizont bursting, such as hemozoin and glycosylphsophatidylinositol (GPI), triggers the febrile response [6]. Since the duration of the IDC varies between different *Plasmodium* species, the periodicity of fever peaks also varies and is a distinctive trait of each species. However, erratic patterns of malaria fever are also commonly observed, possibly due to asynchronous or multiple infections [4, 5, 7]. Malarial fever is only triggered above a certain level of parasitaemia, called the pyrogenic threshold [4, 5, 8]. Therefore, since repeated exposure to malaria results in progressive acquisition of immunity and lower parasite densities, in endemic areas malarial fever (as well as other clinical symptoms) is mainly observed in children [9].

Temperatures above the optimal growth temperature of an organism lead to the accumulation of misfolded and aggregated proteins, which can cause cell damage and death. Therefore, all organisms need mechanisms in place to protect themselves from heat-mediated damage. The universal mechanism to restore proteostasis at higher-than-optimal temperatures, common to both prokaryotes and eukaryotes, is called the heat-shock (HS) response (HSR). It mainly consists in a rapid increase in the expression of chaperone-encoding genes to prevent and revert heat-induced proteome damage [10, 11]. In most eukaryotes, from yeast to humans, the HSR is regulated by the conserved transcription factor HSF1 [12–14]. However, HSF1 is absent from malaria parasites, which are regularly exposed to febrile temperatures and therefore must have a HSR.

We recently described a *P. falciparum* transcription factor of the ApiAP2 family, AP2-HS, which triggers the malarial HSR [15]. While several malarial transcription factors regulate developmental progression or transitions [16], AP2-HS is the only one known to drive a directed transcriptional response to an environmental condition [5, 15]. Although the sequence and structure of AP2-HS is unrelated with HSF1, it plays an analogous role in the regulation of the protective HSR. Three genes are rapidly upregulated in response to febrile temperatures in an AP2-HS-dependent manner: the conserved chaperone-encoding genes *hsp70-1* and *hsp90*, and the gene of unknown function *PF3D7_1421800* [15]. This is an extremely compact HSR compared with the HSF1-dependent HSR described in other eukaryotes, which involves dozens or hundreds of genes [14]. Experiments using recombinant proteins [17] and ChIP-seq [15, 18] showed that AP2-HS recognises the tandem G-box DNA motif in the promoters of the *hsp70-1* and *hsp90* genes, which are direct targets of AP2-HS. In contrast, ChIP-seq experiments revealed that AP2-HS does not bind the *PF3D7_1421800* promoter, which lacks a tandem G-box, indicating that this gene is not a direct target [15]. Besides the AP2-HS-dependent HSR, other transcriptional changes occur during HS in an AP2-HS-independent manner [15, 19, 20], although it is unclear which of these changes are part of a protective response and which ones reflect cell death [5]. Furthermore, several other *P. falciparum* proteins [20–25], as well as protein modifications and metabolites [26,27], are needed for HS survival, but their transcript levels do not appear to increase during HS and therefore they cannot be considered part of a transcriptional “response” [5].

In addition to conferring resistance to thermal stress, AP2-HS may also play a role in resistance to artemisinin and derivatives (ART), the key component of the frontline antimalarial treatment. Parasites deficient for AP2-HS show increased sensitivity to ART, which suggests that the HSR may contribute to ART resistance [15]. Of note, the mode of action of ART involves alkylation of proteins and lipids and formation of reactive oxygen species, which results in general proteome damage, similar to HS, among other consequences for the cell [28, 29]. ART-mediated parasite killing requires activation of the drug by haemoglobin degradation products. Mutations in the K13 protein underlie ART resistance by reducing haemoglobin uptake and drug activation in early ring stage parasites [30–33]. However, functional stress responses are also required for ART resistance, as ART still generates cellular stress in K13 mutants [29]. While the importance of the endoplasmic reticulum (ER)-based unfolded protein response (UPR) for ART resistance is well established [34, 35], whether the other main cellular stress response, the cytoplasm-based HSR, plays a role in ART resistance remains unknown.

Here we set out to characterise the specific conditions that activate the AP2-HS-dependent HSR. We exposed parasites at different stages of their life cycle to different thermal stress conditions, mimicking febrile episodes of different intensity and duration, and assessed the impact on parasite survival and activation of the HSR. We also investigated activation of the AP2-HS-dependent HSR by exposure to ART.

## RESULTS

### Establishment of a new water bath-based standard HS assay

We developed a new standard HS assay to study the effect of febrile temperatures on *P. falciparum in vitro*. We previously used a cell culture incubator-based HS assay [15], but we found that it was difficult to achieve stable, consistent temperatures in an incubator, as others have previously reported [36]. Slight differences in the position of the cultures within the incubator, differences between incubator units, spontaneous temperature fluctuations and the number of times the incubator door was opened during the assay affected the results and compromised reproducibility. Therefore, we developed an alternative assay in which cultures are transferred to a centrifuge tube and HS is performed in a water bath, where temperature is more stable than in an incubator. Furthermore, the temperature of a water bath is easy to control and measure and corresponds well with the temperature of the cultures, whereas the temperature of an incubator does not immediately coincide with the temperature of the cultures within it. Altogether, these characteristics resulted in more consistent and reproducible results using the water bath-based HS assay.

The standard conditions of the new water bath-based HS are 1 h incubation at 41 °C, a temperature and duration similar to the peak of a malaria high fever episode [3–5]. Under these conditions, HS survival at the trophozoite stage was typically ∼70% in wild-type (wt) parasites and ∼10% in parasites with truncated AP2-HS lacking AP2 domain 3 (see below). This is similar to survival of these parasite lines after a 3 h HS at 41.5 °C in an incubator, our previous standard HS conditions [15]. The direct contact with water, which has higher thermal conductivity than air, results in cultures achieving the target temperature faster, explaining why a 1 h at 41 °C HS in a water bath has a similar impact on parasite viability as a longer HS at a higher temperature in an incubator.

### HS duration and temperature conditions needed for activation of the AP2-HS-dependent HSR

We used the water bath-based assay to define how the duration and temperature of a HS affect parasite viability and activation of the HSR. The assays were performed with two 3D7-A subclones: 10E, a subclone with wt AP2-HS, and 10G, a subclone with a spontaneous premature STOP codon in AP2-HS that results in a truncated protein lacking the third AP2 domain (AP2-HStr). 10G does not have any growth defect at 37 °C but is unable to activate the AP2-HS-dependent HSR and has lower HS survival than 10E [15]; therefore, it provides an ideal control to distinguish AP2-HS-dependent from -independent transcriptional changes. We exposed tightly synchronised (5 h age window) cultures of both subclones at the late trophozoite stage [30-35 h post-invasion (hpi)] to a HS of different duration or at different temperature and, at the following cycle, measured parasitaemia by flow cytometry to estimate HS survival. RNA was collected immediately after HS to determine transcript levels of the AP2-HS target genes *hsp70-1* and *hsp90* by reverse transcription (RT)-quantitative PCR (qPCR). For all experiments, control cultures were maintained in parallel in a water bath at 37 °C during HS (**Fig. 1A,D**).

**Fig. 1.**
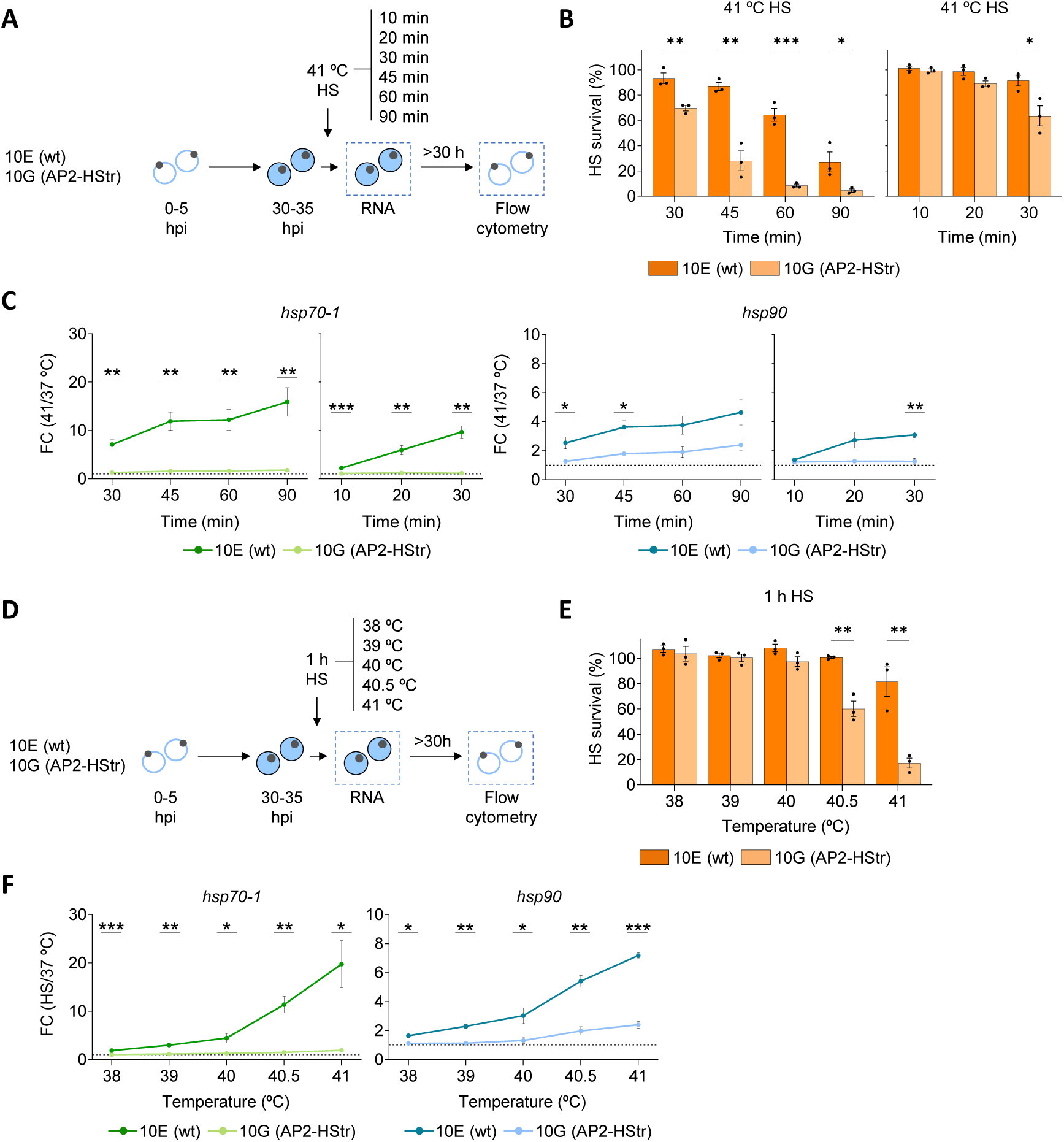
Effect of HS duration and temperature on the activation of the AP2-HS-dependent HSR. **A.** Overview of the experiments to test the effect of HS duration on the activation of the HSR. Tightly-synchronised 10E (wt) and 10G (AP2-HStr) 30-35 hpi (late trophozoite stage) cultures were exposed to a 41 °C HS of variable duration in a water bath. RNA for transcriptional analysis (by RT-qPCR) was collected immediately after HS. To estimate HS survival, parasitaemia was determined by flow cytometry after reinvasion. Control cultures (no HS) were maintained in parallel in a water bath at 37 °C for the duration of the HS. **B.** HS survival after exposing cultures to a 41 °C HS of variable duration, relative to control cultures (no HS). **C.** Fold-change (FC) of normalised *hsp70-1* and *hsp90* transcript levels in cultures exposed to a 41 °C HS of variable duration relative to transcript levels in control cultures (no HS). The horizontal dotted line indicates a FC of 1 (no change). **D.** Overview of the experiments to test the effect of HS temperature on the activation of the HSR, as in panel A but with HS of a constant duration of 1 h and variable temperature. **E.** HS survival after exposing cultures to a 1 h HS at variable temperature, relative to control cultures (no HS). **F.** Fold-change (FC) of normalised *hsp70-1* and *hsp90* transcript levels in cultures exposed to a 1 h HS at variable temperature relative to transcript levels in control cultures (no HS). In all panels, values are the mean ± s.e.m. of n=3 independent biological replicates. Statistically-significant differences between 10E and 10G, calculated using two-sided unpaired Student’s *t*-tests, are indicated by asterisks (*: 0.01<*P*≤0.05; **: 0.001<*P*≤0.01; ***: *P*≤0.001).

Increased duration of HS at 41 °C, from 30 to 90 min, progressively reduced the survival of wt cultures (10E), with essentially 100% survival after a 30 min HS and only ∼27% after a 90 min HS. The viability of cultures expressing truncated AP2-HS (10G) was lower than in 10E for HS of any duration, with more marked differences for HS of 45 min or more (**Fig 1A-B**). Activation of *hsp70-1* and *hsp90*, expressed as the transcript levels fold-change (FC) in cultures exposed to HS relative to control cultures, was observed for HS of any duration (30 to 90 min) and increased with HS duration in 10E, whereas in 10G there was essentially no activation (**Fig. 1C, S1A**). The magnitude of *hsp70-1* activation was larger than that of *hsp90* (maximum activation ∼16-fold vs ∼5-fold, respectively), as previously observed [15], although both genes followed a similar pattern. To confirm these results, we performed experiments using the former standard HS assay (incubator-based HS assay), which revealed a similar pattern of reduced survival and increased activation of the HSR with longer HS (**Fig. S1B-C**).

The observation of a ∼7-fold increase in *hsp70-1* transcript levels after an only 30 min HS led us to investigate the effect of a shorter HS in a separate set of experiments. A ≤20 min HS did not affect survival of either 10E or 10G (**Fig. 1B**) but activated the AP2-HS-dependent HSR in 10E, with a ∼2-fold or ∼6-fold increase in *hsp70-1* transcript levels after HS of only 10 or 20 min, respectively (**Fig. 1B-C, S1A**). Therefore, rapid upregulation of target gene expression, which is a distinctive feature of the HSR in model organisms [10, 14], appears to be conserved in *P. falciparum*. This is, to our knowledge, the fastest transcriptional response described to date in malaria parasites.

Next, we assessed the effect of a 1 h HS at different temperatures on parasite survival and activation of the HSR. At 40 °C or lower temperatures, it did not affect the viability of either 10E or 10G, whereas at higher temperatures up to 41 °C 10G showed a marked decrease in survival and 10E was only moderately affected (**Fig. 1D-E**). The magnitude of the HSR was temperature-dependent, with increased *hsp70-1* and *hsp90* transcript levels observed at temperatures of 38 °C or higher in 10E and only very low-level activation in 10G (**Fig. 1F, Fig. S1D**). Together with the experiments on HS duration, these results show that HS conditions that do not have an impact on parasite viability even in the 10G line (*e.g.*, <30 min at 41 °C or <40.5 °C for 1 h, **Fig. 1B, E**) can activate the HSR.

### The HSR can be activated at most stages of the IDC

To investigate activation of the HSR at different stages of the IDC, tightly synchronised 10E and 10G cultures were exposed to a standard HS (1 h at 41 °C) at four different stages of the IDC (**Fig. 2A-B**). The viability of very early (0-5 hpi) and late (15-20 hpi) ring-stage 10E and 10G cultures was minimally affected by HS, as previously reported [15, 37, 38]. Late trophozoites (30-35 hpi) were the stage most sensitive to HS and showed the largest differences in survival between 10E and 10G, as previously reported [15], whereas schizonts (37-42 hpi) showed survival levels similar to trophozoites in 10E but were more resistant than trophozoites in 10G.

**Fig. 2.**
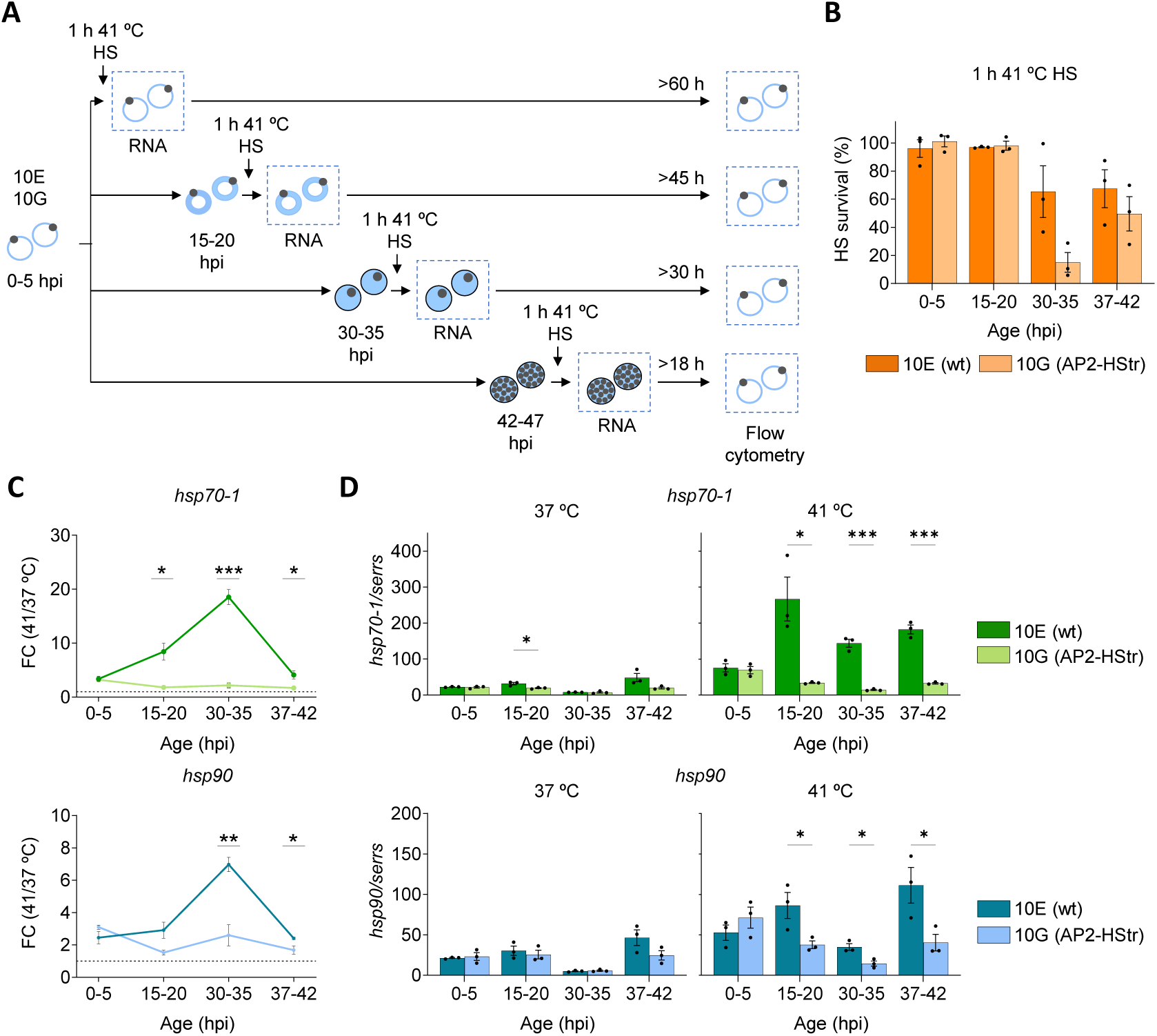
Activation of the AP2-HS-dependent HSR at different stages of the IDC. **A.** Overview of the experiments to test activation of the HSR at different stages of the IDC. Tightly-synchronised 10E (wt) and 10G (AP2-HStr) cultures were exposed to a standard 1 h HS at 41 °C in a water bath at different stages, as indicated. RNA for transcriptional analysis (by RT-qPCR) was collected immediately after HS. To estimate HS survival, parasitaemia was determined by flow cytometry after reinvasion. Control cultures (no HS) were maintained in parallel in a water bath at 37 °C for the duration of the HS. **B.** HS survival after exposing cultures at different stages of the IDC to a 1 h HS at 41 °C, relative to control cultures (no HS). **C.** Fold-change (FC) of normalised *hsp70-1* and *hsp90* transcript levels in cultures at different stages of the IDC exposed to a 1 h HS at 41 °C, relative to transcript levels in control cultures (no HS). **D.** Transcript levels of *hsp70-1* and *hsp90*, normalised against *serine-tRNA ligase* gene (*serrs*) transcripts, in cultures at different stages of the IDC exposed to HS (41 °C) or not (37 °C). In all panels, values are the mean ± s.e.m. of n=3 independent biological replicates. Statistically-significant differences between 10E and 10G, calculated using two-sided unpaired Student’s *t*-tests, are indicated by asterisks (*: 0.01<*P*≤0.05; **: 0.001<*P*≤0.01; ***: *P*≤0.001).

The AP2-HS-dependent HSR was activated at all stages except for very early rings (**Fig. 2C**). In 10E, late trophozoites showed the highest FC in *hsp70-1* and *hsp90* transcript levels between HS-exposed and control cultures. However, *serine-tRNA ligase* (*serrs*)-normalised *hsp70-1* and *hsp90* transcript levels after HS were similar or even higher in late rings and schizonts than in trophozoites (**Fig. 2D**). Therefore, the higher FC in trophozoites than in cultures at other stages is explained by reduced basal expression of *hsp70-1* and *hsp90* at this stage, as part of the cyclic fluctuations in transcript levels of most *P. falciparum* genes along the IDC [39]. The low basal expression of these chaperone-encoding genes in trophozoites may explain the higher sensitivity to HS of parasites at this stage. In very early rings, we detected a ∼3-fold increase in *hsp70-1* and *hsp90* transcript levels after HS in both 10E and 10G (**Fig. 2C-D**), which was confirmed using *ubiquitin-conjugating enzyme* (*uce*) for normalisation (**Fig. S2**). These results suggest that, at this stage, *hsp70-1* and *hsp90* may be activated in response to HS independently from AP2-HS. However, we cannot exclude the possibility that expression of the normalising genes may be altered after HS in very early rings, explaining the apparent increase in *hsp70-1* and *hsp90* transcripts. At other stages, differences between 10E and 10G and genome-wide transcriptomic analysis [15] exclude the possibility that changes in *hsp70-1* and *hsp90* are explained by altered expression of the normalising gene.

### Gametocytes at intermediate or late stages of development do not activate the HSR

We also investigated activation of the HSR in gametocytes at different stages of development [2]. Given that 3D7-A-derived lines, such as 10E and 10G, have a defect in the *gdv1* gene and do not produce gametocytes [40], for these experiments we used different parasite lines. To study the HSR in early gametocytes, we used the 3D7-derived E5 gametocyte-inducible transgenic line (E5ind), in which massive synchronous sexual conversion can be induced by addition of rapamycin [40]. Cultures at stage I or stage II/III of gametocyte development were exposed to a standard HS (1 h at 41 °C) and RNA for transcriptional analysis collected immediately after HS (**Fig. 3A**). Since gametocytes are non-replicative, HS survival was estimated as the gametocytaemia of cultures exposed to HS relative to control cultures, determined by light microscopy analysis of Giemsa-stained smears prepared 2 days after HS. Although these assays showed substantial variability, stage II/III gametocytes, with a mean survival of ∼8%, were clearly more sensitive to HS than stage I gametocytes or any asexual stages (**Fig. 3B-C**, compare with **Fig. 2B**). Furthermore, stage II/III gametocytes showed minimal activation of the expression of *hsp70-1* and *hsp90* after HS, whereas stage I gametocytes activated it at rates comparable to asexual stages (**Fig. 3B, Fig. S3A**). These results suggest a possible causal relationship between inability to activate the HSR and low survival to HS in gametocytes.

**Fig. 3.**
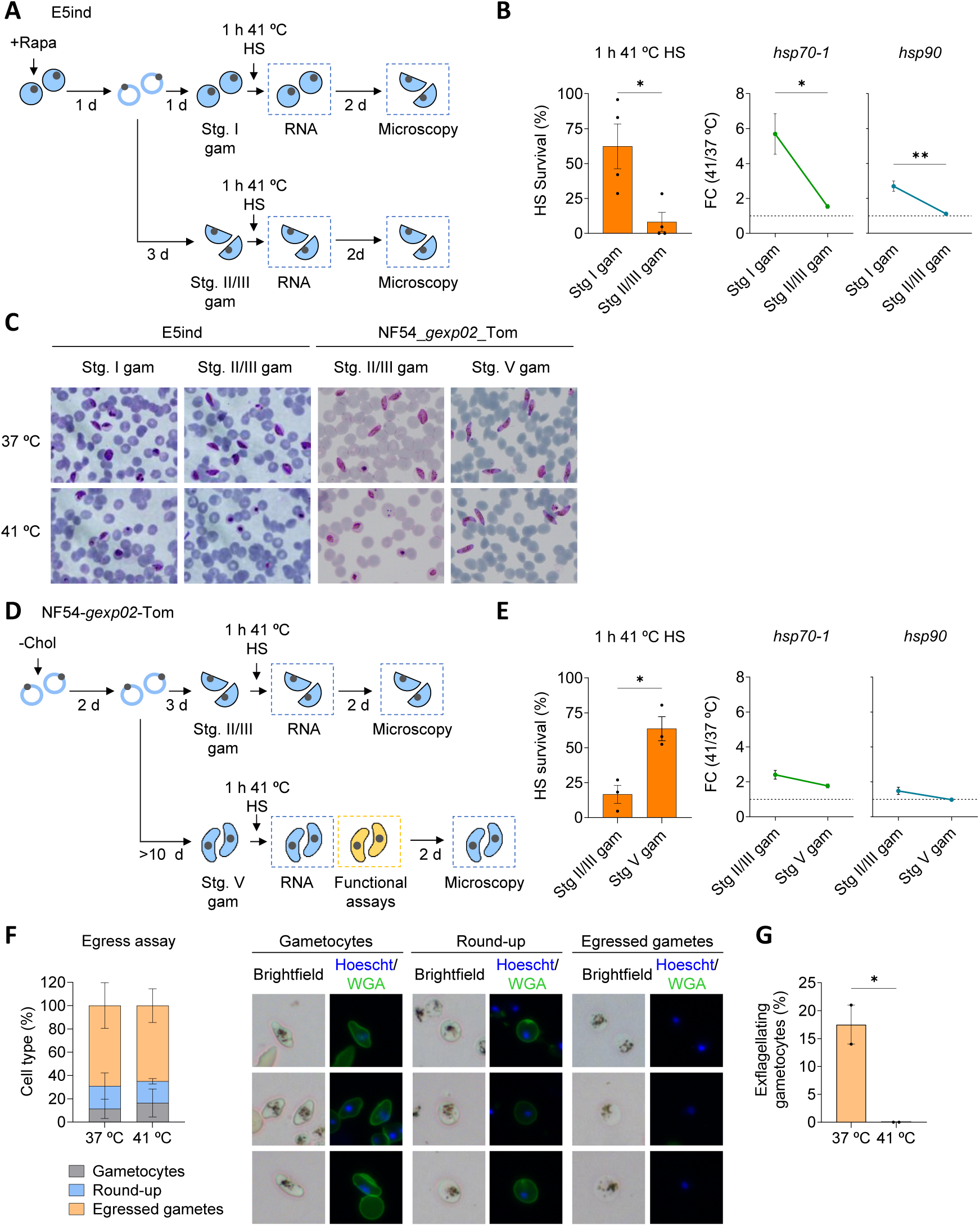
Activation of the AP2-HS-dependent HSR in gametocyes. **A.** Overview of the experiments with the E5ind line to study activation of the HSR at early and intermediate stages of gametocyte development. Sexual conversion was induced by addition of Rapamycin (+Rapa) and cultures were exposed to a standard 1 h HS at 41 °C in a water bath at stage I or stage II/III of gametocyte development. RNA for transcriptional analysis (by RT-qPCR) was collected immediately after HS. To estimate HS survival, gametocytaemia was determined by light microscopy 2 days after HS. Control cultures (no HS) were maintained in parallel in a water bath at 37 °C for the duration of the HS. **B.** HS survival and fold-change (FC) of *hsp70-1* and *hsp90* transcript levels in E5ind gametocyte cultures exposed to a 1 h HS at 41 °C, relative to control cultures (no HS). Values are the mean ± s.e.m. of n=4 independent biological replicates. **C.** Representative images of Giemsa-stained smears prepared 2 days after HS at different stages of gametocyte development, for cultures exposed to HS (41 °C) and their controls (no HS, 37 °C). **D.** Overview of the experiments with the NF54-*gexp02-*Tom line to study activation of the HSR at intermediate and mature stages of gametocyte development. Sexual conversion was induced by choline depletion (- Chol) and cultures were exposed to a standard 1 h HS at 41 °C in a water bath at stage II/III or stage V of gametocyte development. RNA for transcriptional analysis (by RT-qPCR) was collected immediately after HS. To estimate HS survival, gametocytaemia was determined by light microscopy 2 days after HS. Functional assays with mature gametocytes were performed 2.5-5 h after HS. Control cultures (no HS) were maintained in parallel in a water bath at 37 °C for the duration of the HS. **E.** HS survival and fold-change (FC) of *hsp70-1* and *hsp90* transcript levels in NF54-*gexp02-*Tom gametocyte cultures exposed to a 1 h HS at 41 °C, relative to control cultures (no HS). Values are the mean ± s.e.m. of n=3 independent biological replicates. However, for stage V gametocyte survival assays each of the three values is the mean of two HS survival assays performed on aliquots of the same cultures on separate days. **F.** Quantification of gamete activation (round-up) and egress in NF54-*gexp02-*Tom mature (stage V) gametocyte cultures exposed to a 1 h HS at 41 °C and control cultures (37 °C, no HS). The brightfield and fluorescence (blue: Hoechst; green: WGA-Oregon Green 488) images at the right are representative of parasites that were scored as gametocytes (elongated, peripheral WGA signal), round-up (round shape, peripheral WGA signal) or egressed gametes (round shape, no peripheral WGA signal). Values are the mean ± s.e.m. of n=2 independent biological replicates, with ≥200 cells scored for each sample. **G.** Quantification of the percentage of exflagellating gametocytes in NF54-*gexp02-*Tom mature (stage V) gametocyte cultures exposed to a 1 h HS at 41 °C and control cultures (37 °C, no HS). Values are the mean ± s.e.m of n=2 independent biological replicates, with 2 technical replicates for each biological replicate. In all panels, statistically-significant differences between different gametocyte stages (panels B, E) or between HS and no HS (panel G), calculated using two-sided unpaired Student’s *t*-tests (except for panel F, two-way ANOVA), are indicated by asterisks (*: 0.01<*P*≤0.05; **: 0.001<*P*≤0.01; ***: *P*≤0.001).

Since the E5ind line does not produce infective male gametocytes [40] and cultures tend to loose synchronicity after stage III, to study the HSR in mature gametocytes we used the NF54-*gexp02*-Tom reporter line [41]. After inducing sexual conversion by choline depletion [42, 43], NF54-*gexp02*-Tom stage II/III or stage V gametocytes were exposed to a standard HS (1 h at 41 °C) (**Fig. 3D**). Stage II/III gametocytes showed low HS survival (∼17%), similar to E5ind gametocytes at this stage. In contrast, using the same microscopy-based assay, stage V (mature) gametocytes showed higher HS survival (∼64%) (**Fig. 3C-E**). However, transcriptomic analysis using RNA collected immediately after HS showed minimal activation of the HSR in both stage II/III and stage V gametocytes (**Fig. 3E, Fig. S3B**). Together, these results show that, after stage I, gametocytes lose the ability to activate the HSR in response to HS.

While stage V gametocytes, which are a quiescent stage, appeared to be resistant to HS without activation of the HSR, we reasoned that they may be damaged, in spite of normal morphology. To explore this possibility, we performed functional assays 2.5 to 5 h after HS (**Fig. 3D**, **Fig. S3C**). An activation and egress assay showed that HS-exposed stage V gametocytes rounded-up and egressed from their host RBCs at a similar rate as control (no HS) gametocytes (**Fig. 3F**). However, exflagellation assays revealed total absence of exflagellation centres in HS-exposed cultures, in contrast to the ∼17% exflagellation rate observed in control cultures (**Fig. 3G**). This result indicates that at least male mature gametocytes are not viable after HS.

### The gene *PF3D7_1421800* is dispensable for HS survival

*PF3D7_1421800* is a gene of unknown function that, together with *hsp70-1* and *hsp90*, is activated in an AP2-HS-dependent manner in response to HS [15]. However, it does not have a tandem G-box in its promoter and ChIP-seq analysis revealed that it is not a direct target of AP2-HS [15, 18]. Analysis of *PF3D7_1421800* expression in the samples described in **Fig. 1-2** and **Fig. S1-2** confirmed that transcript levels for this gene can increase up to 30 times in 10E cultures exposed to HS (relative to control cultures at 37 °C), but in some experiments a relatively large increase (>10-fold) was also observed in 10G (**Fig. S4**). Activation in response to HS was only observed in trophozoites and schizonts. The activation pattern at different stages, durations and temperatures was clearly distinct from *hsp70-1* and *hsp90*. Of note, the basal normalised transcript levels of this gene are very low at all IDC stages (*e.g.*, ∼300-fold lower than *hsp70-1* in trophozoites). Inspection of expression profiles of this gene in published datasets available in PlasmoDB revealed that the gene is expressed at much higher levels in mature gametocytes than during the IDC, which was confirmed by the analysis of its transcript levels in the gametocyte samples described above (**Fig. 4A**). However, the expression of the gene was not increased in response to HS in stage II/III or V gametocytes.

**Fig. 4.**
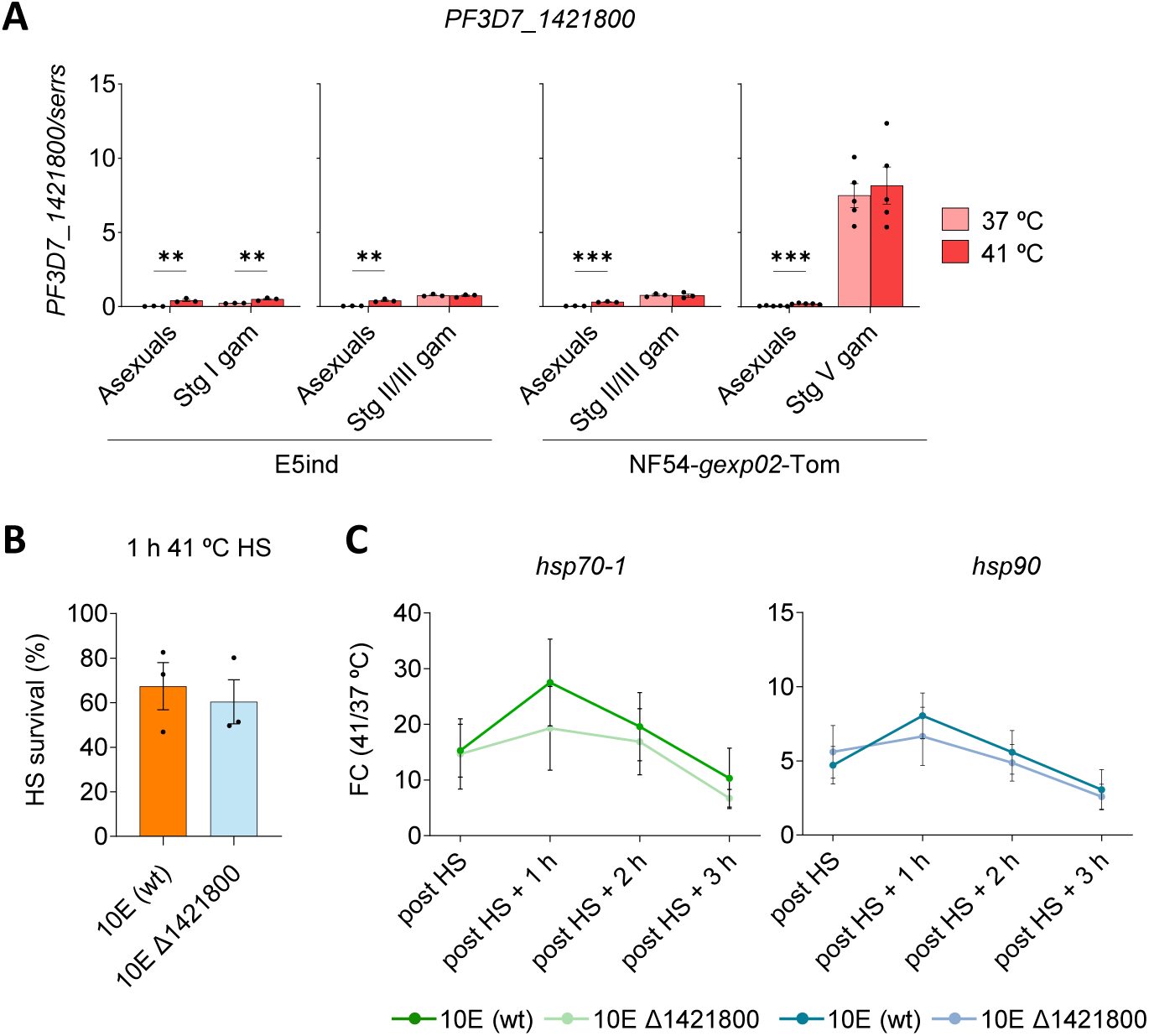
Characterisation of the role of the *PF3D7_1421800* gene in the HSR. **A.** Transcript levels of *PF3D7_1421800*, normalised against the *serine-tRNA ligase* gene (*serrs*), in asexual blood stages at the late trophozoite stage and different stages of gametocyte development, exposed to a standard 1 h at 41 °C HS (41 °C) or not (37 °C). Values are the mean ± s.e.m. of n=3-5 independent biological replicates (see individual data points). **B.** HS survival after exposing 30-35 hpi (late trophozoite stage) 10E (wt) and 10E_Δ1421800 cultures to a standard 1 h HS at 41 °C, relative to control cultures (no HS). Values are the mean ± s.e.m. of n=3 independent biological replicates. **C.** Fold-change (FC) of *hsp70-1* and *hsp90* transcript levels in 10E (wt) and 10E_Δ1421800 cultures immediately after HS (post HS) or after additional incubation at 37°C following HS, relative to transcript levels in control cultures (no HS). Values are the mean ± s.e.m. of n=3 independent biological replicates. In all panels, statistically-significant differences between HS and no-HS cultures (panel A) or between 10E and 10E_Δ1421800 (panels B-C), calculated using two-sided unpaired Student’s *t*-tests, are indicated by asterisks (*: 0.01<*P*≤0.05; **: 0.001<*P*≤0.01; ***: *P*≤0.001).

To further investigate the role of this gene in the HSR, we generated a *PF3D7_1421800* knockout (KO) line (10E_Δ1421800) by transfection of the 10E line (**Fig. S5A**). The KO line had a multiplication rate, HS survival and activation of *hsp70-1* and *hsp90* expression in response to HS similar to the wt parental line (**Fig. 4B-C, Fig. S5B-C**). These results indicate that this gene is dispensable for normal progression through the IDC and for the activation of the AP2-HS-dependent protective HSR.

Finally, we explored a possible role of this gene in switching off the HSR. It is well established that activation of the HSR is transient, in *P. falciparum* [15] as well as in other organisms [10, 14], possibly because prolonged activation is detrimental for the cells [44]. We speculated that *PF3D7_1421800* may be involved in the downregulation of the HSR, which may explain its activation after HS only in parasites able to activate the HSR, in spite of not being an AP2-HS direct target. However, the *hsp70-1* and *hsp90* transcript levels dynamics at 1 to 3 h after HS showed no differences between wt and 10E_Δ1421800 cultures, excluding a role for this gene in the deactivation of the HSR (**Fig. 4C, Fig. S5C**).

Together, these results suggest that *PF3D7_1421800* is a gametocyte-specific gene that only has residual expression during the IDC. Its intriguing activation in response to HS in trophozoites and schizonts is of unclear significance, and not fully dependent on full-length AP2-HS.

### A sublethal dose of dihydroartemisinin (DHA) triggers the AP2-HS-dependent HSR

In a previous study, several parasite lines in which the entire *ap2-hs* gene was disrupted (Δ*ap2-hs* lines) showed higher susceptibility than their wt controls to DHA, the active metabolite of ART, both at the ring and at the trophozoite stages. In contrast, the 10G parasite line, expressing truncated AP2-HS lacking the third AP2 domain (AP2-HStr), showed only a very minor increase in sensitivity to DHA at the trophozoite stage [15]. These results may suggest that the HSR, which is not activated in either Δ*ap2-hs* or 10G, contributes to DHA survival. However, the higher sensitivity to DHA of Δ*ap2-hs* lines compared to 10G raise an alternative possibility: Δ*ap2-hs* lines may suffer from constitutive proteome damage, even in the absence of stress, as a consequence of low basal expression of *hsp70-1* and *hsp90* [15]. This could make them more sensitive to additional proteotoxic damage (*e.g.*, by DHA). This latter possibility is supported by the observation of growth defects at 37 °C in Δ*ap2-hs* lines but not in 10G, which expresses normal basal levels of *hsp70-1* and *hsp90* [15].

To investigate if DHA activates the HSR, we measured *hsp70-1, hsp90* and *PF3D7_1421800* transcript levels in 30-35 hpi 10E and 10G cultures exposed to sublethal doses of DHA (3 h pulse). Survival was similar between 10E and 10G at all the DHA concentrations tested (**Fig. 5A**). However, activation of *hsp70-1* and *hsp90* in response to DHA was evident in 10E (up to 10-fold increase) but not in 10G (**Fig. 5B, Fig. S6**), whereas *PF3D7_1421800* was only modestly activated in both parasite lines. This result shows that exposure to DHA activates the AP2-HS-dependent HSR, providing evidence for activation of the HSR by proteotoxic conditions different from thermal stress. However, activation of the HSR did not result in substantially increased survival to DHA.

**Fig. 5.**
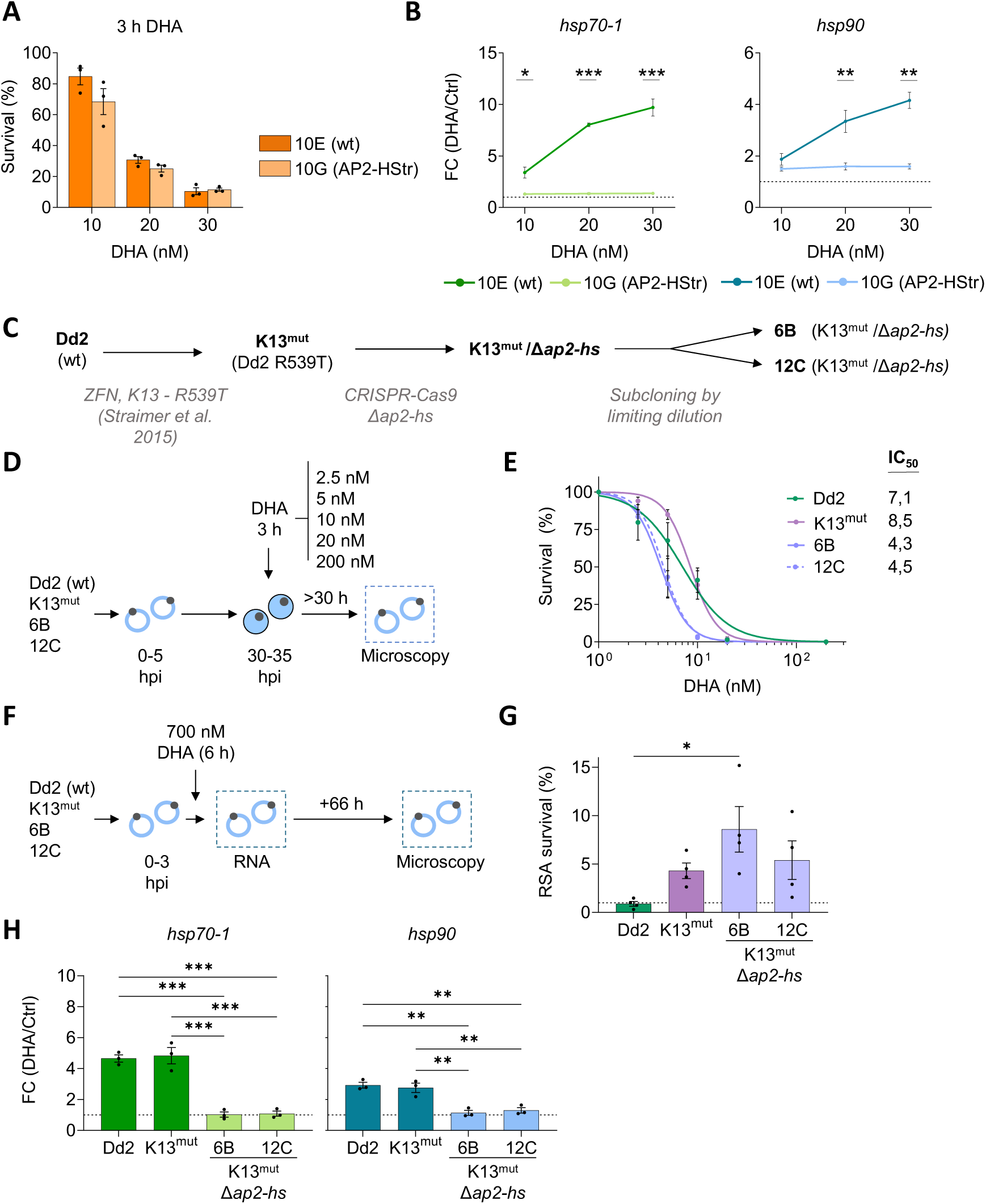
DHA triggers the AP2-HS-dependent HSR. **A.** Survival of 10E (wt) and 10G (AP2-HStr) 30-35 hpi (late trophozoite stage) cultures after exposure to a 3 h DHA pulse at different concentrations, relative to controls not exposed to DHA. **B.** Fold-change (FC) of *hsp70-1* and *hsp90* transcript levels in cultures exposed to DHA at different concentrations relative to transcript levels in control cultures (no DHA). In panels A and B, values are the mean ± s.e.m. of n=3 independent biological replicates. Statistically-significant differences between 10E and 10G, calculated using two-sided unpaired Student’s *t*-tests, are indicated by asterisks (*: 0.01<*P*≤0.05; **: 0.001<*P*≤0.01; ***: *P*≤0.001). **C.** Schematic of the parasite lines used for the experiments in the following panels. **D.** Overview of the experiments to determine the IC_50_ of K13 and AP2-HS mutants to DHA. Tightly-synchronised 30-35 hpi cultures (late trophozoite stage) were exposed to a 3 h DHA pulse at different concentrations. To estimate survival, parasitaemia was determined by light microscopy after reinvasion. **E.** Survival of 30-35 hpi cultures after exposure to a 3 h DHA pulse at different concentrations, relative to controls not exposed to DHA. Values are the mean ± s.e.m. of n=2 independent biological replicates. IC_50_ values for each parasite line were calculated using dose-response sigmoidal curves. **F.** Overview of the ring survival assay (RSA) experiments with K13 and AP2-HS mutants. For the RSA, tightly-synchronised 0-3 hpi cultures were exposed to a 6 h 700 nM DHA pulse. RNA for transcriptional analysis (by RT-qPCR) was collected immediately after the pulse. To estimate survival, parasitaemia was determined by light microscopy after reinvasion. **G.** Survival of the different parasite lines in the RSA. Survival above 1% (dotted horizontal line) is considered an ART-resistant phenotype. Values are the mean ± s.e.m. of n=4 independent biological replicates. **H.** Fold-change (FC) of *hsp70-1* and *hsp90* transcript levels in cultures exposed to DHA relative to control cultures (no DHA) in the RSA. Values are the mean ± s.e.m. of n=3 independent biological replicates. In panels G and H, statistically-significant differences between parasite lines, calculated using one-way ANOVA, are indicated by asterisks (*: 0.01<*P*≤0.05; **: 0.001<*P*≤0.01; ***: *P*≤0.001).

### Activation of the AP2-HS-dependent HSR after a high-dose DHA pulse

Having established that a sublethal DHA pulse triggers the AP2-HS-dependent HSR, we set out to investigate if a clinically-relevant dose of DHA also triggers this response. Since all wt parasites exposed to such a dose of DHA die, we performed these experiments using ART-resistant K13 mutants [30–33]. Additionally, with these experiments we aimed to determine if the AP2-HS driven HSR is needed for ART resistance in K13 mutants.

We generated the double transgenic line K13^mut^/Δ*ap2-hs*, in which the entire *ap2-hs* gene was deleted in the previously described Dd2 R539T (hereafter K13^mut^) line [32]. This parasite line, of Dd2 genetic background, had been engineered to introduce an ART-resistance mutation in the K13 protein. Subclones 6B and 12C of K13^mut^/Δ*ap2-hs* were used for all the experiments (**Fig. 5C, Fig. S7A**). For consistency with previous studies, K13^mut^/Δ*ap2-hs* subclones and their controls were regularly maintained at 35 °C. As observed for other Δ*ap2-hs* lines [15], the K13^mut^/Δ*ap2-hs* subclones showed a reduced multiplication rate at either 35 °C or 37 °C, were highly sensitive to HS and failed to activate the expression of *hsp70-1*, *hsp90* and *PF3D7_1421800* in response to HS (**Fig. S7B-D**). The K13^mut^ line and the parental wt Dd2 line had a similar IC_50_ for DHA at the trophozoite stage (30-35 hpi), consistent with previous reports showing that ART resistance does not result in changes in the IC_50_ for this drug at the trophozoite stage [45]. In contrast, K13^mut^/Δ*ap2-hs* subclones had a lower IC_50_ (**Fig. 5D-E**), consistent with previous reports for Δ*ap2-hs* lines [15].

Next, we analysed K13^mut^/Δ*ap2-hs* and control lines using the ring-survival assay (RSA), the gold standard *in vitro* assay to predict ART resistance *in vivo* [45]. In this assay, tightly synchronised cultures at the very early ring stage (0-3 hpi) are exposed to a 6 h DHA pulse at a clinically-relevant concentration (700 nM) and parasitaemia is measured by light microscopy 66 h later to determine the proportion of surviving parasites. While the Dd2 wt line showed a sensitive phenotype (<1% survival), the K13^mut^ and the K13^mut^/Δ*ap2-hs* lines (6B and 12C subclones) showed a resistant phenotype (>1% survival), indicating that the AP2-HS-dependent HSR is not essential for ART resistance in K13 mutants (**Fig. 5F-G**). Transcriptional analysis of RNA samples collected immediately after the 6 h DHA pulse showed a clear increase in *hsp70-1* and *hsp90* transcripts in the wt Dd2 and the K13^mut^ lines, but not in the K13^mut^/Δ*ap2-hs* subclones (**Fig. 5H, Fig. S8**). This result demonstrates that a DHA pulse mimicking a clinical ART treatment triggers the AP2-HS-dependent HSR, but this is not needed for survival of K13 mutants.

## DISCUSSION

We systematically investigated which conditions activate the *P. falciparum* AP2-HS-dependent HSR, including thermal stress of different duration and intensity and chemical stress induced by the antimalarial drug DHA. We also investigated which parasite blood stages can activate the HSR, including different asexual stages and gametocytes, the sexual forms that mediate transmission to mosquitoes. We found that both severe and mild thermal stress conditions, including conditions that have essentially no impact on parasite viability, can activate the HSR. Treatment of asexual parasites with DHA also activated the HSR, indicating that the parasite may use the AP2-HS-based response to withstand non-thermal types of proteome-damaging stress, although in the specific case of DHA, activation of the HSR did not substantially increase survival. The HSR can be activated at all stages of the IDC, with the exception of very young rings. In contrast to IDC stages, gametocytes beyond stage I of development did not activate the HSR. Strikingly, intermediate-stage and mature gametocytes exposed to a HS mimicking a malaria severe fever peak were not viable, which raises the intriguing possibility that malaria patients experiencing high fever episodes may transiently become non-infective to mosquitoes.

Our new water bath-based HS assay, together with the use of a control parasite line in which the AP2-HS-dependent HSR is not activated, enabled the sensitive detection of low-level HSR activation with high confidence. We found that a HS of only 10 minutes can lead to low magnitude but consistent activation of the HSR, demonstrating that the malarial HSR is a rapid response, similar to all other eukaryotes in which it has been investigated [10, 14]. Likewise, HS at a temperature as low as 38 °C also led to low level but consistent HSR activation. The conditions that activated the HSR fall well within the range of thermal conditions encountered by the parasite during natural malaria fever episodes [3–5], suggesting that the parasite frequently activates the HSR during clinical malaria infections. Furthermore, axillary temperature, which is regularly used in epidemiological studies to measure fever, typically gives values up to 1 °C below the internal body temperature to which the parasite is actually exposed [46–48], implying that essentially all malaria fevers involve conditions likely to activate the HSR. The observation that activation of the HSR occurs under HS duration and temperature conditions that, even in parasites unable to activate the AP2-HS-dependent HSR, do not have an impact on parasite viability, suggests that *P. falciparum* responds to thermal stress in a ‘preventive’ manner. Activating the HSR early during a fever episode, before it actually produces irreversible damage and compromises viability, can provide a survival advantage to the parasite: preventing the formation of misfolded or aggregated proteins, rather than repairing them after they accumulate, is a more effective strategy to survive thermal stress.

Our results confirm previous reports indicating that ring stages are intrinsically resistant to febrile temperatures [15, 37, 38], even in a parasite line in which the AP2-HS-dependent HSR cannot be activated. Late trophozoites are the most sensitive stage, and schizonts show intermediate levels of sensitivity. The level of HS resistance somehow parallels the basal levels of *hsp70-1* and *hsp90* along the IDC, with high expression in rings and the lowest expression in late trophozoites [39, 49, 50], a pattern that we confirmed. Therefore, the variability in HS sensitivity between IDC stages may be determined by the basal levels of these key chaperones. Despite the changes in basal transcript levels of chaperone-encoding genes during the IDC, we demonstrate that the AP2-HS-dependent HSR can be activated at all stages of the IDC except for very early rings. Of note, the HSR can be activated in 15-20 hpi ring stages, in which the response appears to be unnecessary for survival, which supports the idea of ‘preventive’ activation of the HSR under non-lethal conditions. However, it is possible that the viability of ring stages would be affected by a more severe HS, as previously reported [19, 38], and therefore the capacity to activate the HSR response in rings may provide a survival advantage.

In contrast to the capacity of most IDC stages to activate the AP2-HS-dependent HSR, experiments with gametocytes revealed that only stage I gametocytes can activate it, whereas more mature sexual stages cannot. This was paralleled by low resistance to HS of gametocytes at intermediate or mature stages, revealed either by microscopy analysis of parasites 2 days after HS or by functional assays. These unexpected observations raise the intriguing possibility that, in natural malaria infections, gametocytes may be killed by malarial high fever episodes. In the case of immature gametocytes (*e.g.*, stage II/III), which develop in the bone marrow [2], we cannot exclude the possibility that this niche confers some level of physical protection against temperature changes. However, mature (stage V) gametocytes are found in the peripheral circulation and therefore are fully exposed to febrile temperatures. Even though stage V gametocytes exposed to HS can be activated to gametes and egress from RBCs, we show that males are unable to exflagellate. Thus, since both functional males and females are needed for successful transmission, we predict that malaria patients suffering from severe fever episodes become temporarily non-infectious to mosquitoes. In contrast, their future infectivity may be increased because febrile temperatures enhance sexual conversion rates [43], resulting in a complex, double-edged effect of fever on transmission. Some published field-based studies support the idea that febrile patients have reduced infectivity for both *P. falciparum* [51–53] and *P. vivax* [54]. Future studies specifically designed to address this question should confirm whether high fever results in transient non-infectiousness and establish if there is a temperature threshold above which malaria patients become non-infectious.

We demonstrate that the AP2-HS-depenendent HSR is activated in response to DHA treatment. However, in contrast to the ER-based UPR, which is needed for ART resistance [28, 29, 34, 35], activation of the AP2-HS-dependent HSR does not appear to be essential for ART resistance, as DHA sensitivity was similar between wt parasites and parasites expressing truncated AP2-HS, which cannot activate the HSR. Furthermore, survival of K13 mutants in the clinically-relevant RSA [45] is similar between *ap2-hs* wt and KO lines. Therefore, the AP2-HS-dependent HSR is activated by both thermal stress and DHA-induced proteotoxic stress, but only in the case of thermal stress activation of the response is associated with increased survival. A possible explanation for this observation is that DHA exerts pleiotropic effects on the parasite, such that the drug affects not only proteome integrity but also disrupts lipids, generates reactive oxygen species and interferes with multiple cellular processes through protein alkylation [28, 29]. While the upregulation of *hsp70-1* and *hsp90* in response to DHA likely contributes to counteracting proteome damage, it may not protect from the additional damage on other cell components caused by DHA. Thus, the reduced survival of Δ*ap2-hs* lines to DHA sublethal pulses [15] most likely reflects basal proteome damage, which leaves parasites at the edge of proteostasis collapse, rather than inability to activate the HSR.

## MATERIALS & METHODS

### Parasite cultures

The 3D7-A subclones 10E and 10G [55], the gametocyte-inducible line E5ind [40], the NF54-*gexp02*-Tom reporter line [41], and the Dd2 and K13^mut^ (Dd2 R539T) lines [32] have been previously described. Parasites were cultured under standard conditions with either B^+^ or O^+^ RBCs at 3% haematocrit in a 5% CO_2,_ 2% O_2_, balance N_2_ atmosphere. The 10E, 10G, E5ind and 10E_Δ1421800 lines were regularly cultured at 37 °C under static conditions in standard RPMI-based medium supplemented with Albumax II (Invitrogen). The K13^mut^/Δ*ap2-hs* line (6B and 12C subclones) and its controls (Dd2 and K13^mut^ lines) were maintained under the same conditions but at 35°C. The NF54-*gexp02*-Tom line was cultured at 37°C under shaking conditions with the same culture medium supplemented with 2 mM choline chloride (Sigma-Aldrich C7527).

### Generation of transgenic parasite lines

To disrupt the *PF3D7_1421800* gene (10E_Δ1421800 line), we used the CRISPR-Cas9 system (**Fig. S5A**). Cloning was performed using the InFusion system (Takara). The donor plasmid (pL6 _HRs1421800_hdhfr) was generated from a modified pL6-egfp-yfcu plasmid [31] in which the *yfcu* cassette was previously removed [40]. The homology regions (HRs) were PCR-amplified from genomic DNA (HR1: positions -460 to 0 from the start codon of *PF3D7_1421800*; HR2, positions +1 to +439 from the stop codon) and cloned into SpeI/AflII and EcoRI/AatII sites, respectively, replacing the GFP5’ region (HR1) or the GFP3’ region and sgRNA cassette (HR2) in the original plasmid. The Cas9 plasmid (pDC2_sh_sgRNA1421800) was derived from the pDC2-Cas9-HDHFRyFCU plasmid [56] after removing the *hdhfr-yfcu* cassette by digestion with EcoRI followed by treatment with T4 DNA polymerase (NEB, n. M0203S) to generate blunt ends, digestion with HpaI and religation with T4 ligase (Roche, n. 10481220001). The sgRNA targeting the *PF3D7_1421800* coding region was prepared by annealing forward and reverse oligonucleotides and cloned into BpiI-digested plasmid using the InFusion system (Takara, n. 639642). Ring-stage 10E cultures were transfected with 15 µg of PvuI-linearized donor plasmid and 60 µg of circular Cas9 plasmid using a BioRad GenePulser Xcell electroporator. After transfection, cultures were selected for 4 days with 10 nM WR99210 (Jacobus Pharmaceuticals), and fresh RBCs were provided to the culture once per week until parasites were observed 15 days after transfection. Diagnostic PCR analysis of genomic DNA was used to confirm correct edition and absence of detectable parasites with wt *PF3D7_1421800* locus (**Fig. S5A**). All primers used for PCR amplification of HRs, generation of sgRNAs and diagnostic PCR are described in **Table S1**.

To knockout the *ap2-hs* gene from the K13^mut^ line using the CRISPR/Cas9 system (K13^mut^/Δ*ap2-hs* line), we used the same plasmids and approach previously described [15] (**Fig. S7A**), with a modification in the quantity of plasmids used for transfection. In brief, ring stage cultures were transfected with 30 µg of circular pDC2_wo/hdhfr_ap2hs_sgRNA3’ plasmid, which encodes Cas9 and one sgRNA, and 60 µg of PvuI-linearized pL7-ap2hs_KO_sgRNA5’ donor plasmid, which contains the HRs flanking a *hdhfr* expression cassette and encodes a second sgRNA. After transfection using a BioRad GenePulser Xcell electroporator, selection with WR99210 and maintenance were performed as described above, and parasites were observed 13 days after transfection. Subclones were obtained by limiting dilution. Correct edition was confirmed by diagnostic PCR analysis of genomic DNA (**Fig. S7A**) with primers described in **Table S1**.

### HS assays

For the standard HS assays, cultures were synchronised to a 5 h age window by Percoll purification of late stages followed by sorbitol lysis (to remove late stages) 5 h later [57]. At 30-35 hpi, 4 ml of each culture at ∼1.5% parasitemia were transferred to 15 ml centrifuge tubes and gassed for 5 s with regular parasite culture gas mixture (5% CO_2,_ 2% O_2_, balance N_2)_. The tubes were placed in a water bath at 41°C (HS samples) or at 37 °C (control samples; 35 °C for experiments with K13^mut^/Δ*ap2-hs* lines). The temperature of the water bath was calibrated using a precision glass thermometer and validated at each experiment. The tubes were placed in a plastic rack inside the water bath, avoiding the use of glass beakers to hold the tubes (we found that the temperature of the water inside a beaker was lower than in the rest of the water bath). After 1h in the corresponding water baths, cultures were resuspended (typically by pipetting up and down) and two 100 µl aliquots of each sample were transferred to a 96-well plate and cultured for >30 h to assess HS survival by flow cytometry (at this time, all viable parasites had completed the IDC and reinvaded, even if HS induced delays in life cycle progression). The remaining volume of each sample (∼4 ml) was centrifuged at 1,500 rpm for 5 min and the pellet resuspended in 20 pellet volumes (12 for rings) of TRIzol reagent (Invitrogen, n. 15596018) and stored at -80 °C. To measure parasitaemia by flow cytometry, 5 µl of culture were resuspended in 800 µl of PBS, stained with 1 µl of SYTO11 nucleic acid stain (Invitrogen, n. S7573) and incubated for 1 min before analysis in a BD FACSCalibur flow cytometer (BD Biosciences), as previously described [58]. The parasitaemia of cultures exposed to HS relative to the parasitaemia of control cultures maintained in parallel at 37 °C (or 35 °C) was used to determine HS survival. For the incubator-based standard HS assay, 30-35 hpi cultures were exposed for 3 h to a 41.5°C HS in a cell culture incubator, within an air-tight incubation chamber, as previously described [15].

For the HS duration experiments, one tube of each parasite line was placed in the water bath for each HS duration and, after the selected incubation time, each sample was taken from the water bath and processed as mentioned above. For the HS temperature experiments, six different water baths at different temperature were used to simultaneously expose identical aliquots of the same synchronised cultures to the different temperatures. To assess the effect of HS at different stages of the IDC, HS was applied to cultures at different points of the IDC derived from the same tightly synchronised culture.

### RNA extraction and RT-qPCR

RNA was collected in TRIzol immediately after HS or DHA treatment and purified using a method suitable for low amounts of RNA [59]. In brief, RNA was purified using the RNeasy Mini Kit (Qiagen), DNase treated using the RNase-free DNase set (Qiagen, n. 79254) and cDNA prepared by reverse transcription using the AMV Reverse Transcription Kit (Promega, n. A3500), as previously described [57]. The resulting cDNAs were used to quantify transcript abundance by qPCR, using the PowerSYBR Green master mix (Applied Biosystems) and the standard curve method [57]. Each sample was tested in triplicate and negative controls (reactions without reverse transcriptase) tested in duplicate for a selection of the genes analysed. Transcript levels were normalised using *serine-tRNA ligase* (*serrs*, *PF3D7_0717700*) or *ubiquitin-conjugating enzyme* (*uce*, *PF3D7_0812600*). Primers used for qPCR are described in **Table S1**.

### Gametocyte experiments

To induce sexual conversion in the E5ind line, cultures were sorbitol-synchronised and 20 h later treated with rapamycin for 1h to induce DiCre-mediated recombination [40]. The next day, cultures were placed in complete medium supplemented with 10% B^+^ human serum instead of Albumax II, to support gametocyte development, and heparin (Merck, H3149-50KU) was added to remove asexual parasites (20 U/ml) [60]. The culture medium (with human serum and heparin) was changed daily throughout the experiment and, in some experiments, heparin was removed the day before the assays. A standard HS was performed on day 2 (stage I) or on day 4 (stage II/III) after rapamycin treatment. To determine HS survival, 2 days after HS the gametocytaemia of cultures exposed to HS and control cultures maintained in parallel at 37 °C was measured by light microscopy analysis of Giemsa-stained smears (pyknotic parasites not counted). In all experiments, in cultures exposed to HS we observed reduced gametocytemia (relative to controls), rather than major alterations in gametocyte stage composition. This indicates that HS resulted in death and disintegration or collapse of sensitive gametocytes, rather than only developmental arrest.

To induce sexual conversion in the NF54-*gexp02*-Tom line, cultures were sorbitol-synchronised and maintained in medium without choline for 48 h under static conditions [41]. Choline depletion induces sexual conversion [42]. After 48 h, Albumax II-supplemented medium was replaced by medium supplemented with B^+^ human serum or with Albumax II plus 1x ITS-X (Gibco, n. 51500056), which is a recently described human serum alternative to support full development of viable gametocytes [61]. While initial experiments were performed with human serum-supplemented medium, we found that ITS-X was much easier to obtain and prepare and improved gametocyte development, so we used it for the remaining experiments. We did not observe differences in the results of the experiments between cultures supplemented with human serum or with ITS-X. At 48 h after choline depletion, we also added heparin to the culture medium to remove asexual forms. Medium was changed daily for the next 4 days and then every 2 or 3 days until the end of the experiment. NF54-*gexp02*-Tom gametocyte cultures were exposed to a standard HS at day 3 after adding heparin (stage II/III of sexual development) or between days 11 and 16, when the vast majority of gametocytes had reached full maturity (stage V). HS survival was determined as described above for the E5ind line, by light microscopy analysis of Giemsa-stained smears prepared 2 days after HS.

Exflagellation assays [62] were performed with mature gametocyte cultures 2.5-5 h after HS, typically on day 14 after adding heparin. 30 µl of cultures exposed to HS and their controls were placed in 1.5 ml tubes at 37°C. Cultures were spun and 10 µl of the supernatant removed. Next, gamete activation was induced by diluting 1:2 with ookinete media containing 100 µM xanthurenic acid (XA) and incubating at room temperature. For each sample, ∼10 µl were loaded to a haemocytometer and, 12 min after inducing activation, exflagellation centres were counted by light microscopy with a 40X objective. Each culture was tested twice (two technical replicates). To calculate the percentage of gametocytes that exflagellated, we used the number of exflagellation centres per µl determined in the haemocytometer and the number of gametocytes per µl that was estimated from the number of RBCs per µl (determined in the same haemocytometer) and the gametocytaemia of the culture, determined from Giemsa-stained smears prepared immediately before the assay.

Gamete activation and egress assays were performed with mature gametocyte cultures 2.5-5h after HS, typically at day 15 after adding heparin, following a previously described protocol [63]. After Percoll purification to enrich for gametocytes, parasites from cultures exposed to HS or control cultures were resuspended in 150 µl of complete culture medium (Albumax II and ITS-X-supplemented) and stained with WGA-Oregon Green 488 conjugate (Invitrogen, n. W6748) at a final concentration of 2.5 µg/ml and Hoechst 33342 (Sigma-Aldrich, n. TA9H97BAECD2) at a final concentration of 2 µg/ml. After 15 min at 37°C, samples were washed with RPMI-HEPES (at the same concentration as in the parasite culture medium), followed by resuspension in culture medium with 20 µM XA and ≥15 min incubation at room temperature to induce gamete activation and egress. After spinning and resuspending in PBS, wet preparations were inspected in a fluorescence microscope. Parasites were scored as gametocytes (elongated and with peripheral WGA-488 signal), activated gametes (rounded-up but not egressed from RBCs, *i.e.*, with peripheral WGA-488 signal) or activated and egressed gametes (absence of WGA-488 signal).

### DHA sensitivity assays

Cultures were synchronised to a 5 h age window using the Percoll-sorbitol method, as described above, and adjusted to a 1% (experiments with Dd2-derived lines) or 1.5% (experiments with 10E and 10G) parasitaemia. At 30-35 hpi, cultures were exposed to different concentrations of DHA for 3 h. For 10E and 10G experiments, immediately after removing DHA, ∼4 ml of each sample was collected in TRIzol for RNA extraction (as described above). For the rest of the culture, after washing once with RPMI-HEPES, samples were resuspended in complete parasite culture medium and 100 µl of each sample were transferred in triplicate to a 96-well plate and placed under culture again. At the following cycle, ∼60-65 h after synchronisation, Giemsa-stained smears were prepared for each sample in triplicate. Parasitaemias were determined by light microscopy inspection of the smears and parasite survival calculated relative to the parasitaemia in the control (untreated) culture. For IC_50_ determinations, drug concentrations were log_10_-transformed, and percent survival values were fit to a dose-response sigmoidal curve.

### Ring Survival Assay (RSA)

RSAs were performed according to a standard protocol [45]. Cultures of the Dd2, K13^mut^ and K13^mut^/Δ*ap2-hs* (6B and 12C subclones) parasite lines were synchronised to a 3 h age window using the Percoll-sorbitol method and adjusted to 1% parasitaemia. Immediately after synchronisation, at 0-3 hpi, a fraction of each culture (1.6 ml) was exposed to either a physiological dose of DHA (700 nM final concentration) or dimethyl sulfoxide (DMSO) carrier (1% final concentration) as a control, and incubated for 6 h at 35 °C. Next, 1 ml of each culture was collected, resuspended in TRIzol and stored at -80 °C for future RNA extraction. DHA was removed from the remaining culture and, after washing with RPMI-HEPES, 100 µl of each culture were transferred in triplicate to a 96-well plate and cultured for 66 h at 35 °C. At this time, Giemsa-stained smears were prepared for each sample in duplicate/triplicate and parasitaemias determined by light microscopy.

### Statistical analysis

Statistical analysis was performed using GraphPad Prism 10.6.1. To determine the statistical significance of differences between parasite lines, stages or conditions, we used two-tailed unpaired Student’s *t*-test for pairwise comparisons and, for multiple comparisons, unpaired one-way ANOVA with Tukey’s correction for multiple testing.

## ACKNOWLEDGMENTS

We are grateful to Dr. Pablo Suárez-Cortés (Max Planck Institute for Infection Biology, Berlin, Germany) for providing protocols and useful advice for functional assays with gametocytes. The following reagents were obtained through BEI Resources, NIAID, NIH: *Plasmodium falciparum*, Strain Dd2, MRA-156, contributed by Thomas E. Wellems; and *Plasmodium falciparum*, Strain Dd2_R539T, MRA-1255, contributed by David A. Fidock. This work was supported by grants PID2019-107232RB-I00 and PID2022-137863OB-I00 to A.C. from the Spanish Ministry of Science and Innovation (MCIN)/ Agencia Estatal de Investigación (AEI, 10.13039/501100011033), co-funded by the European Regional Development Fund (ERDF, European Union). N.R. was supported by a fellowship from MCIN/AEI/10.13039/501100011033 (PRE2020-094497). This research is part of ISGlobal’s Program on the Molecular Mechanisms of Malaria, which is partially supported by the Fundación Ramón Areces. We acknowledge support from the grant CEX2023-0001290-S funded by MCIN/AEI/10.13039/501100011033, and support from the Generalitat de Catalunya through the CERCA Program.

## Notes

### Competing Interest Statement

The authors have declared no competing interest.

