## Supplementary Figs. and Table for "Thermal and non-thermal stress conditions activate the *Plasmodium falciparum* AP2-HS-dependent heat-shock response"

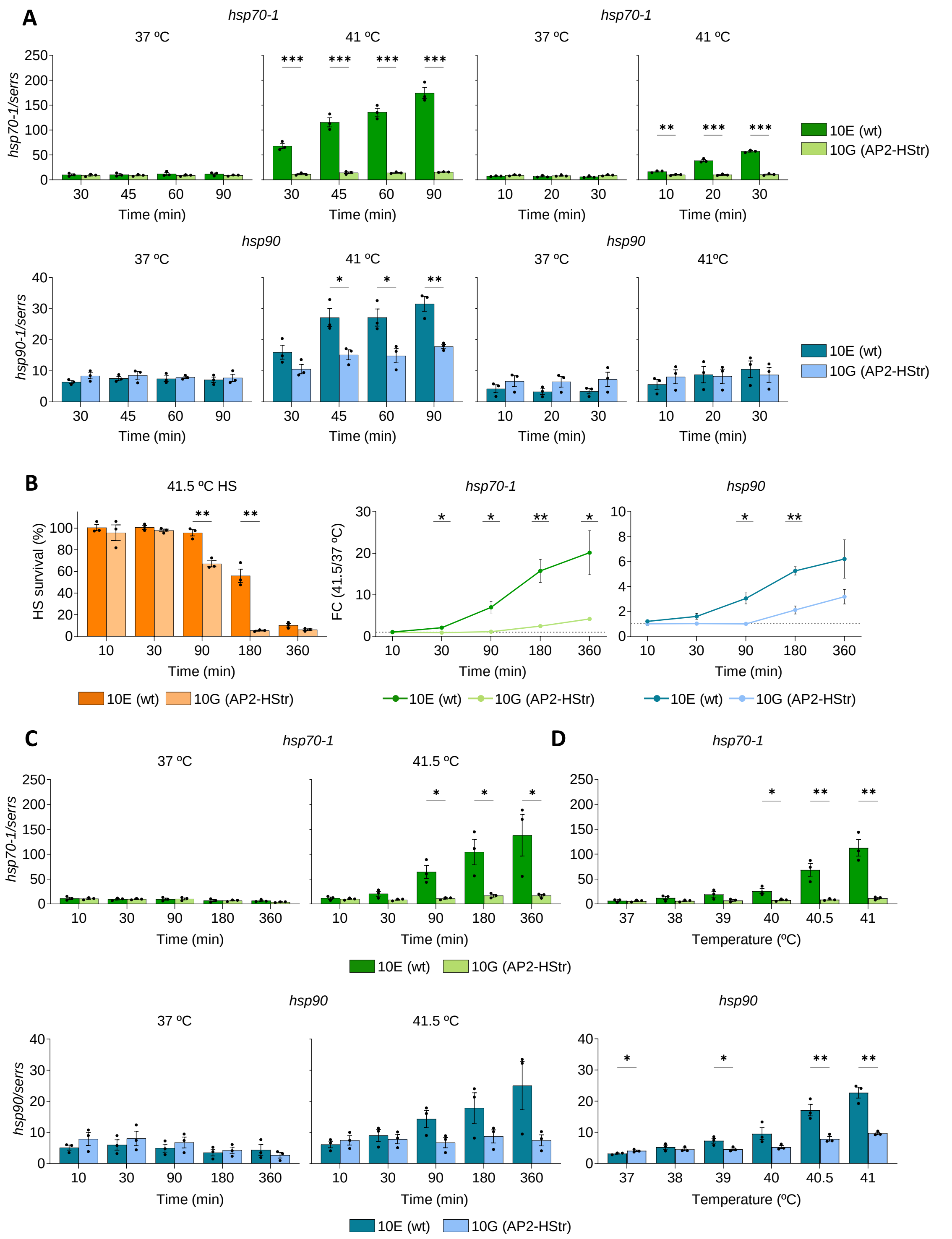


**Fig. S1. Transcript levels of *hsp70-1* and *hsp90* after HS of different duration or at different temperatures. A.** Transcript levels of *hsp70-1* and *hsp90*, normalised against *serine-tRNA ligase* (*serrs*) transcripts, in cultures exposed to a HS of variable duration (41 ºC) or not (37 ºC) in a water bath. **B.** Left, HS survival after exposing cultures to a 41.5 ºC HS of variable duration in an incubator, relative to control cultures (no HS). Right, fold-change (FC) of normalised *hsp70-*

*1* and *hsp90* transcript levels in cultures exposed to a 41.5 ºC HS of variable duration relative to transcript levels in control cultures (no HS). The horizontal dotted line indicates a FC of 1 (no change). **C.** Transcript levels of *hsp70-1* and *hsp90*, normalised against *serrs* transcripts, in cultures exposed to a HS (41.5 ºC) of variable duration in an incubator or not (37 ºC). **D.** Transcript levels of *hsp70-1* and *hsp90*, normalised against *serrs* transcripts, in cultures exposed to a 1 h HS at variable temperature in a water bath or control cultures (37 ºC). In all panels, values are the mean ± s.e.m. of n=3 independent biological replicates. Statistically-significant differences between 10E and 10G, calculated using two-sided unpaired Student's *t*-tests, are indicated by asterisks (*: 0.01<*P*≤0.05; **: 0.001<*P*≤0.01; ***: *P*≤0.001).

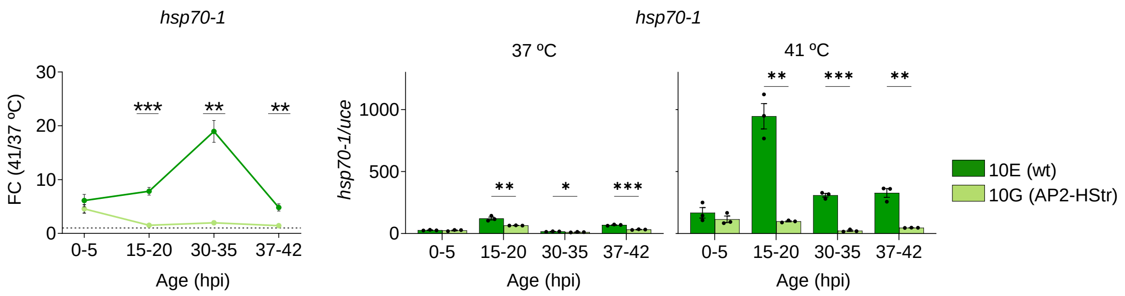


**Fig. S2. Transcriptional changes after HS at different stages of the IDC using *uce* as a normalising gene.** Left, fold-change (FC) of *ubiquitin-conjugating enzyme* (*uce*)-normalised *hsp70-1* transcript levels in cultures at different stages of the IDC exposed to a 1 h HS at 41 ºC, relative to transcript levels in control cultures (no HS). Right, *uce*-normalised *hsp70-1* transcript levels in cultures at different stages of the IDC exposed to HS (41 ºC) or not (37 ºC). In all panels, values are the mean ± s.e.m. of n=3 independent biological replicates. Statistically-significant differences between 10E and 10G, calculated using two-sided unpaired Student's *t*-tests, are indicated by asterisks (*: 0.01<*P*≤0.05; **: 0.001<*P*≤0.01; ***: *P*≤0.001).

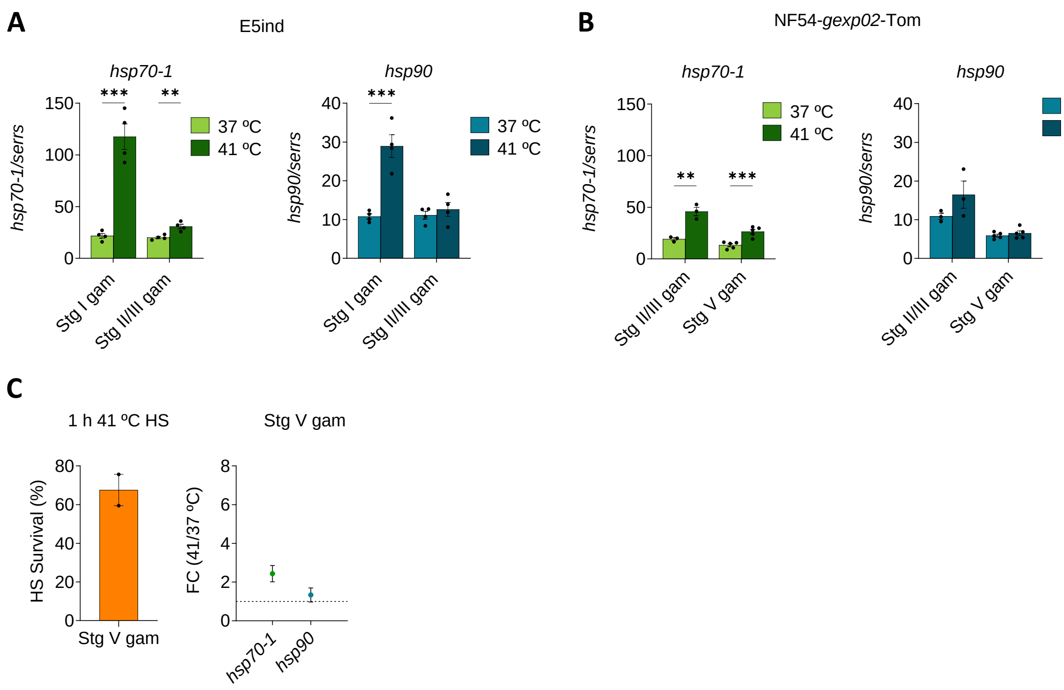


**Fig. S3. Transcript levels of *hsp70-1* and *hsp90* after HS in gametocytes. A.** Transcript levels of *hsp70-1* and *hsp90*, normalised against *serine-tRNA ligase* (*serrs*) transcripts, in E5ind cultures at different stages of gametocyte development exposed to a 1 h HS (41 ºC) or not (37 ºC). Values are the mean ± s.e.m. of n=4 independent biological replicates. **B.** Transcript levels of *hsp70-1* and *hsp90*, normalised against *serrs* transcripts, in NF54-*gexp02*-Tom cultures at different stages of gametocyte development exposed to a 1 h HS (41 ºC) or not (37 ºC). Values are the mean ± s.e.m. of n=3 (stage II/III gametocytes) or 5 (stage V gametocytes) independent biological replicates. In panels A and B, statistically-significant differences between HS and no-HS cultures, calculated using two-sided unpaired Student's *t*-tests, are indicated by asterisks (*: 0.01<*P*≤0.05; **: 0.001<*P*≤0.01; ***: *P*≤0.001). **C.** HS survival and fold-change (FC) of normalised *hsp70*-1 and *hsp90* transcript levels in NF54-*gexp02*-Tom gametocyte cultures exposed to a 1h HS at 41ºC, relative to control cultures (no HS). These data correspond to the gametocyte cultures used for egress and exflagellation assays and is analogous to experiments presented in main Fig. 3E. Values are the mean ± s.e.m of n=2 independent biological replicates, with HS survival data corresponding to two technical replicates (independent experiments performed on the same cultures) for each biological replicate.

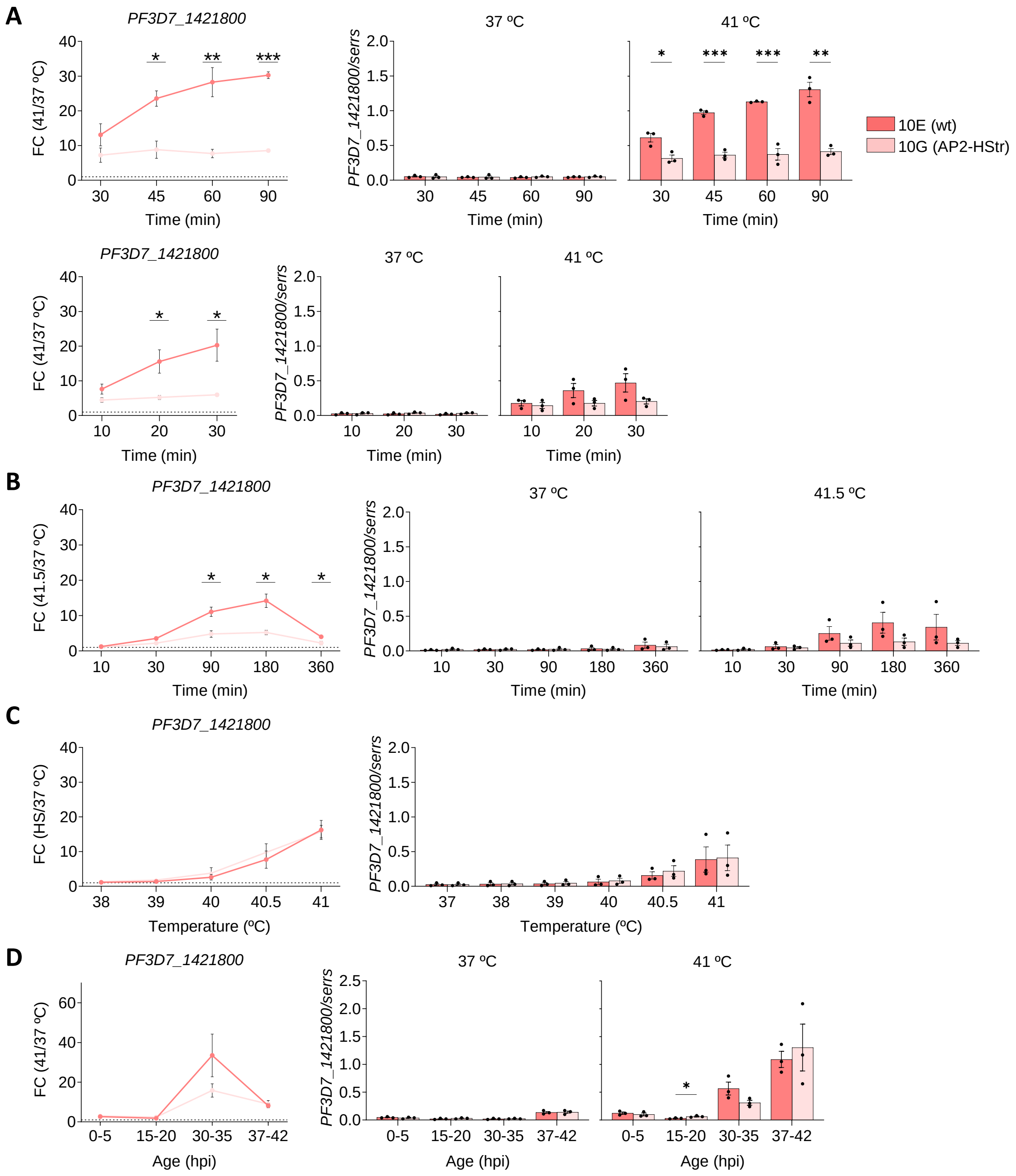


**Fig. S4. Changes in *Pf3D7_1421800* transcript levels after HS with different conditions. A.** Left, fold-change (FC) of *Pf3D7_1421800* normalised transcript levels in 10E (wt) and 10G (AP2-HStr) cultures exposed to a 41 ºC HS of variable duration in a water bath relative to transcript levels in control cultures (no HS). The horizontal dotted line indicates a FC of 1 (no change). Right, transcript levels of *PF3D7_14241800*, normalised against *serine-tRNA ligase* (*serrs*) transcripts*,* in cultures exposed to a HS of variable duration (41 ºC) or not (37 ºC). **B.** Same as in panel A, for experiments performed in an incubator. **C.** Same as in panel A, for experiments with a 1 h HS at variable temperature. **D.** Same as in panel A, for experiments in which cultures were exposed to a 1 h HS at 41 ºC at different stages of the IDC. In all panels, values are the mean ± s.e.m. of n=3 independent biological replicates. Statistically-significant differences between 10E and 10G, calculated using two-sided unpaired Student's *t*-tests, are indicated by asterisks (*: 0.01<*P*≤0.05; **: 0.001<*P*≤0.01; ***: *P*≤0.001).

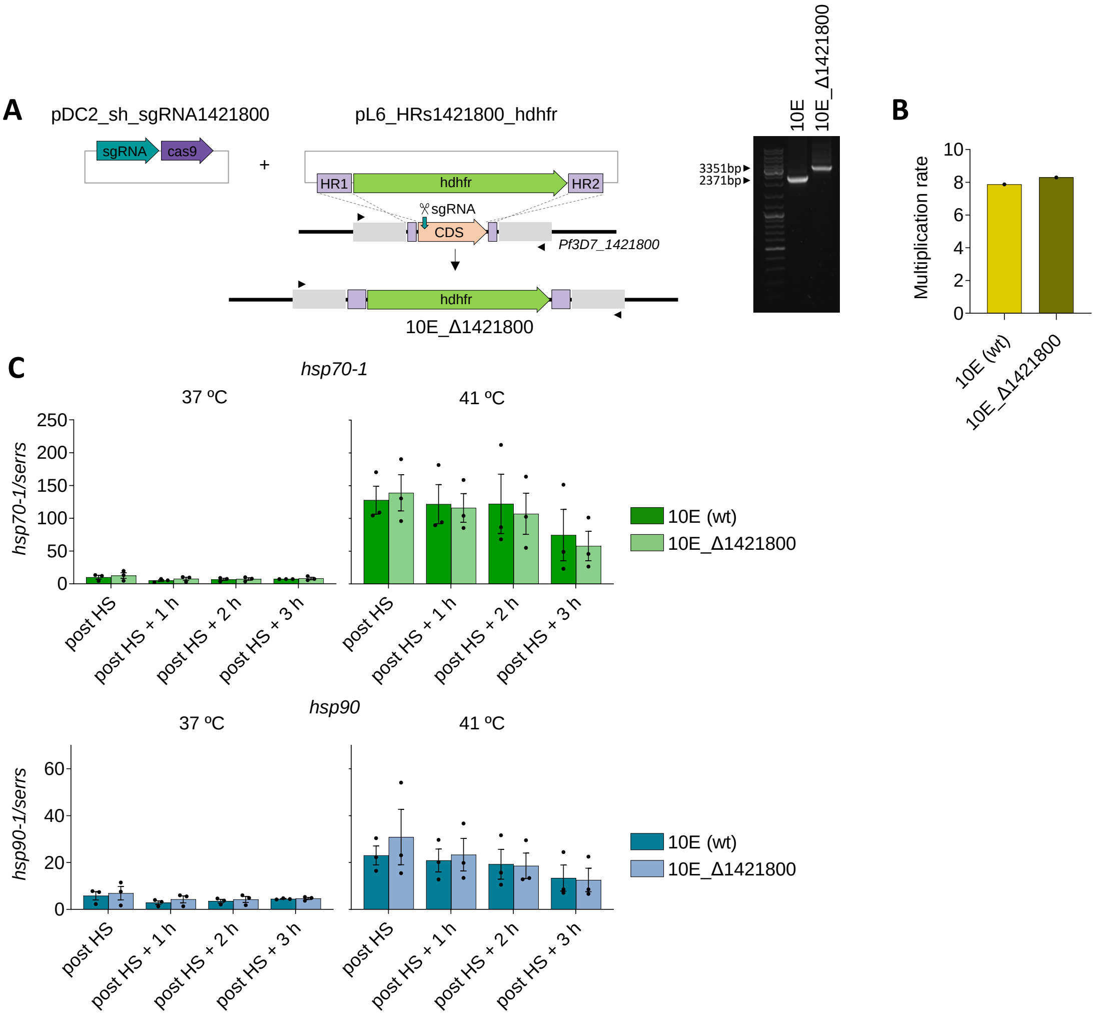


**Fig. S5. Generation and characterisation of the *PF3D7_1421800* KO line. A.** Schematic of the CRISPR-Cas9 strategy to knockout the *PF3D7_1421800* gene and diagnostic PCR analysis of the *PF3D7_1421800* locus showing correct editing. PCR was performed with primers (arrowheads) external to the homology regions (HRs) (Table 1, primers 12/13). The position targeted for cleavage by the guide RNA (sgRNA) is indicated. **B.** Multiplication rate of 10E and 10E_Δ1421800 at 35 ºC and 37 ºC, determined by flow-cytometry measures of parasitaemia at two consecutive cycles. Data from n=1 independent biological replicate. **C.** Transcript levels of *hsp70-1* and *hsp90* normalised against *serine-tRNA ligase*

(*serrs*) transcripts, in 10E (wt) and 10E_Δ1421800 cultures immediately after a standard 1 h HS at 41 ºC (post HS) or after additional incubation at 37ºC. Values are the mean ± s.e.m. of n=3 independent biological replicates. No statistically significant differences were observed at any time point between 10E and 10E_Δ1421800, using a two-sided unpaired Student's *t*-test.

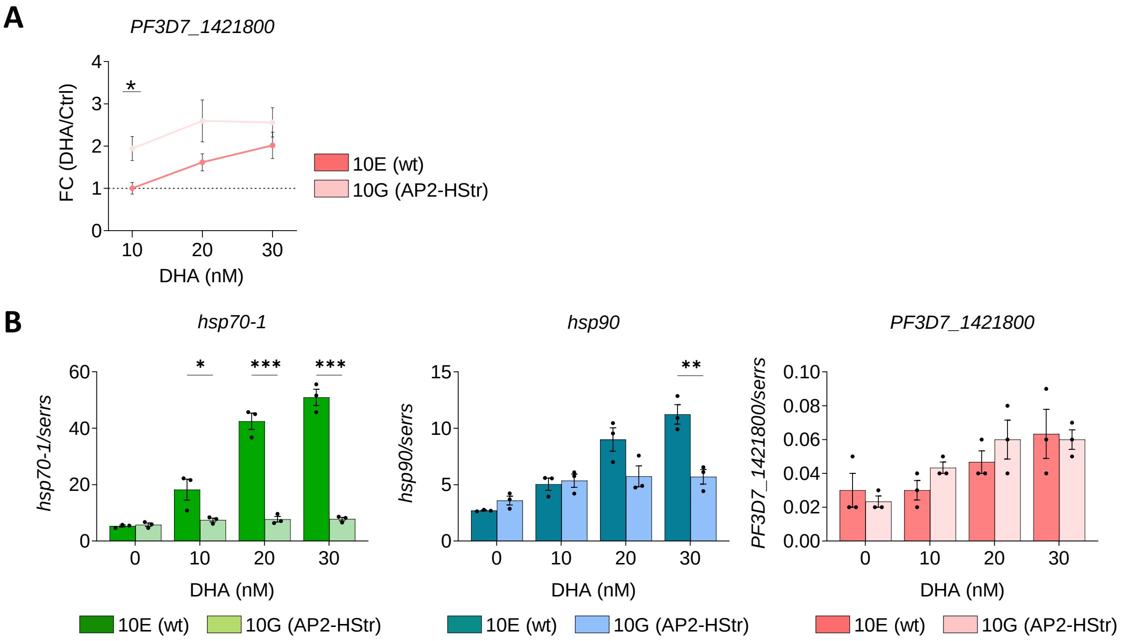


**Fig. S6. Transcriptional changes after exposure to a DHA pulse. A.** Fold-change (FC) of normalized *PF3D7_1421800* transcript levels in 10E (wt) and 10G (AP2-HStr) cultures exposed to a 3 h DHA pulse at different concentrations, relative to controls not exposed to DHA. The horizontal dotted line indicates a FC of 1 (no change). **B.** Transcript levels of *hsp70-1*, *hsp90*, and *PF3D7_1421800*, normalised against *serine-tRNA ligase* (*serrs*) transcripts, in 10E (wt) and 10G (AP2-HStr) cultures exposed to a 3 h DHA pulse at different concentrations. In all panels, values are the mean ± s.e.m. of n=3 independent biological replicates. Statistically-significant differences, calculated using two-sided unpaired Student's *t*-tests, are indicated by asterisks (*: 0.01<*P*≤0.05; **: 0.001<*P*≤0.01; ***: *P*≤0.001).

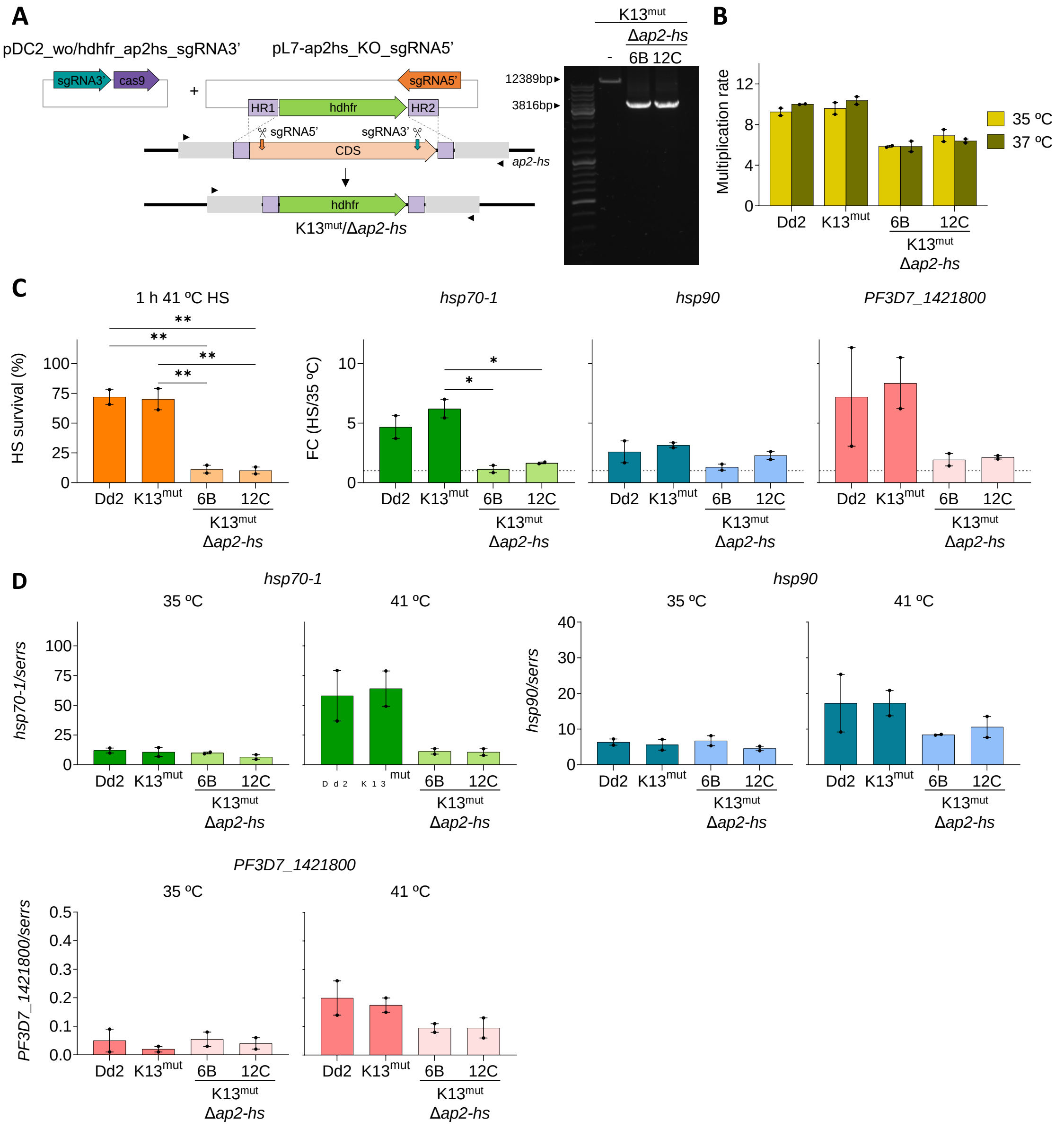


**Fig. S7. Generation and characterisation of K13 and AP2-HS mutant lines.** **A.** Schematic of the CRISPR-Cas9 strategy to knockout the *ap2-hs* gene and diagnostic PCR analysis of the *ap2-hs* locus to validate correct editing. PCR was performed with primers (arrowheads) external to the homology regions (HRs) (Table 1, primers 1/2). The positions targeted for cleavage by the guide RNAs (sgRNAs) are indicated. **B.** Multiplication rate of Dd2 (wt), K13 and AP2-HS mutants at 35 ºC and 37 ºC, determined by flow-cytometry measures of parasitaemia at two consecutive cycles. Values are mean ± s.e.m. of n=2 independent biological replicates. **C.** Left, HS survival after exposing cultures of Dd2 (wt) and mutant parasite lines at the late trophozoite stage (30-35 hpi) to a 1 h HS at 41 ºC, relative to control cultures not exposed to HS. Right, fold-change

(FC) of normalised *hsp70-1*, *hsp90,* and *PF3D7_1421800* transcript levels in wt and mutant cultures exposed to HS relative to transcript levels in control cultures (no HS). **D.** Transcript levels of *hsp70-1*, *hsp90* and *PF3D7_1421800*, normalised against *serine-tRNA ligase* (*serrs*) transcripts, in cultures exposed to HS (41 ºC) or not (35 ºC). In all panels, values are the mean ± s.e.m. of n=3 independent biological replicates. Statistically-significant differences between strains, calculated using one-way ANOVA, are indicated by asterisks (*: 0.01<*P*≤0.05; **: 0.001<*P*≤0.01; ***: *P*≤0.001).

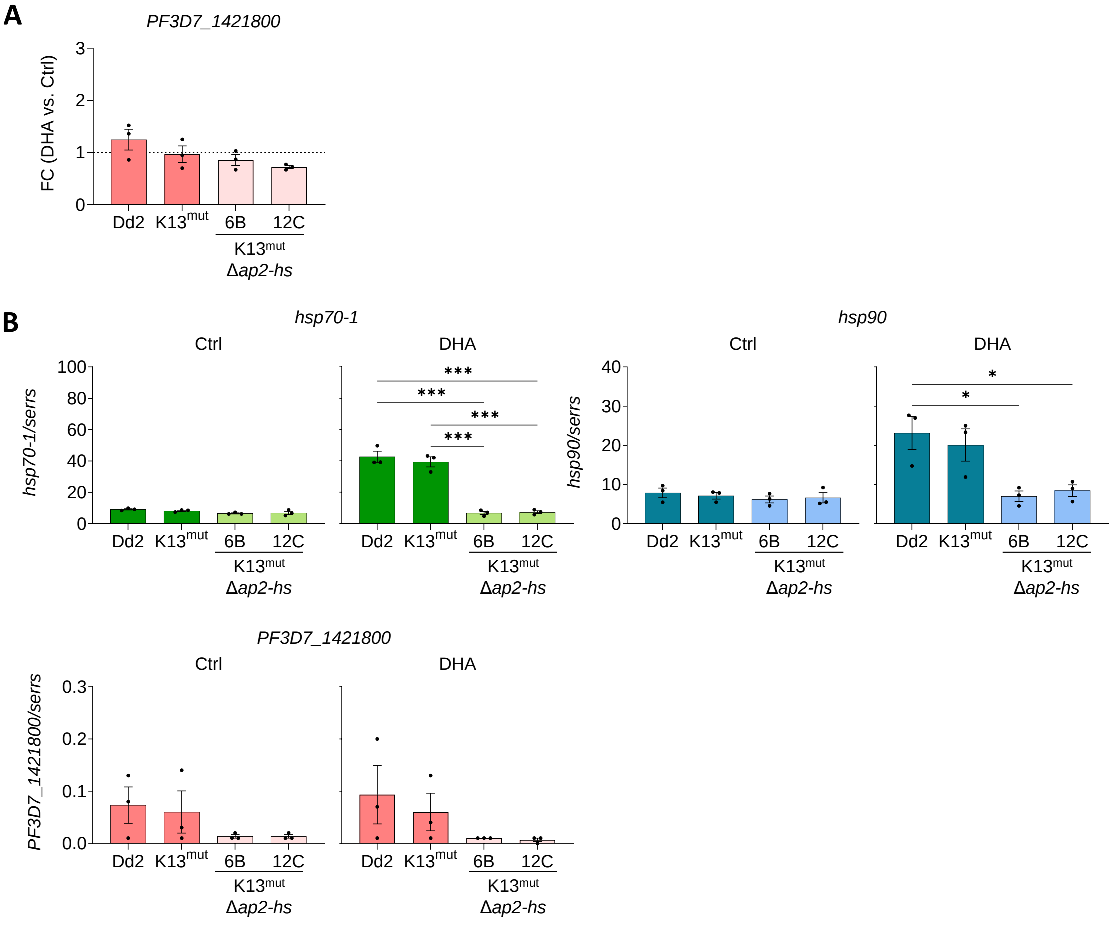


**Fig. S8. Transcriptional changes in K13 and AP2-HS mutant lines in the ring survival assay (RSA). A.** Fold-change (FC) of normalized *PF3D7_1421800* transcript levels in Dd2 wt and mutant cultures exposed to a 6 h 700 nM DHA pulse at 0-3 hpi in the RSA, relative to transcript levels in control cultures (no DHA). **B.** Transcript levels of *hsp70-1*, *hsp90*, and *PF3D7_1421800*, normalised against *serine-tRNA ligase* (*serrs*) transcripts, in cultures exposed to a DHA pulse in the RSA or not (Ctrl). In all panels, values are the mean ± s.e.m. of n=3 independent biological replicates. Statistically-significant differences between parasite lines, calculated using one-way ANOVA, are indicated by asterisks (*: 0.01<*P*≤0.05; **: 0.001<*P*≤0.01; ***: *P*≤0.001).

| | Name | Sequence (5' → 3') | Use |
|---|---|---|---|
| 1 | ap2hs_-94_Fw | CAGTTGATGATTACATCTCTG | Ext. PCR *ap2hs* KO Fw |
| 2 | ap2hs_end_+718_Rv | TTCATCACTTGTTAAGCATCC | Ext. PCR *ap2hs* KO Rv |
| 3 | ap2hs_+10399_Fw | AGAAATGAACAAAACATATTGGG | Copy nº qPCR *ap2hs* KO Fw |
| 4 | ap2hs_+10510_Rv | TTATATTTGTTATAGGTTCCTCC | Copy nº qPCR *ap2hs* KO Rv |
| 5 | 1421800_-460_Fw_InF | gccggggaggactagTATTGTGTACTTTTCCAATGAAG | Amp. HR1 1421800 Fw |
| 6 | 1421800_+2_Rv | TCCTTCCTCTACTTTTACATTC | Amp. ext HR1 1421800 Rv |
| 7 | 1421800_-25_Rv_InF | ttacaaaatgcttaagATTGTATTTTTATTTTATAATTTAA | Amp. HR1 1421800 Rv |
| 8 | 1421800_+1116_Fw_InF | attaaatctagaattcTTTAATGAAGATACTACTACTTACA | Amp. HR2 1421800 Fw |
| 9 | 1421800_+1555_Rv_InF | gaaaagtgccacctgacgtcAACATGTTTGCCTTAATACAATAAA | Amp. HR2 1421800 Rv |
| 10 | 1421800_+23_sgRNA_Fw | taagtatataatattACATCCTTATAATAGTTACGgttttagagctagaa | sgRNA 1421800 Fw |
| 11 | 1421800_+43_sgRNA_Rv | ttctagctctaaaacCGTAACTATTATAAGGATGTaatattatatactta | sgRNA 1421800 Rv |
| 12 | 1421800_-588_Fw | CAAGTCTGTCCTACATACATAC | Ext. PCR 1421800 KO |
| 13 | 1421800_+1783_Rv | CCCTATACAAATGTGTTTGAATT | Ext. PCR 1421800 KO |
| 14 | Seryl_+590_Fw | AAGTAGCAGGTCATCGTGGTT | qPCR seryl Fw |
| 15 | Seryl_+747_Rv | TTCGGCACATTCTTCCATAA | qPCR seryl Rv |
| 16 | Uce_+67_Fw | GGTGTTAGTGGCTCACCAATAGGA | qPCR uce Fw |
| 17 | Uce_+155_Rv | GTACCACCTTCCCATGGAGTA | qPCR uce Rv |
| 18 | hsp70-1_+1155_Fw | TGCAGCTGTACAAGCAGCC | qPCR hsp70-1 Fw |
| 19 | hsp70-1_+1315_Rv | GACTCTTTTTAGCAGGTATGG | qPCR hsp70-1 Rv |
| 20 | hsp86_+1860_Fw | ATCAGAATTTGGATGGTCCGC | qPCR hsp90 Fw |
| 21 | hsp86_+1989_Rv | TGATATAATTGGGTGACGAGC | qPCR hsp90 Rv |
| 22 | PF3D7_1421800_+217_Fw | GTTGATGATAATGTGAAAAACCG | qPCR 1421800 Fw |
| 23 | PF3D7_1421800_+350_Rv | TATAACGTGTGAAATTCAATATCC | qPCR 1421800 Fw |
| 24 | K13_+368_Fw | GCAAATCTTATAAATGATGATTCTGG | Amp. K13 locus Fw* |
| 25 | K13_+2471_Rv | GCTAATAAGTAATATCAATATAAGGG | Amp. K13 locus Rv* |
| 26 | K13_+1311_Fw | GGTATTAAATTTTTACCATTCCCATTAGTATTTTGTATAGG | Sanger seq. K13 mut Fw* |
| 27 | K13_+1937_Rv | TGTTCATTATCAATACCTCCAAC | Sanger seq. K13 mut Rv |

*From Straimer et al. 2015, PMID 25502314

**Table S1. Primers used in this study.**
